# ISG15 is required for the dissemination of *Vaccinia virus* extracellular virions

**DOI:** 10.1101/2022.10.27.514002

**Authors:** Martina Bécares, Manuel Albert, Celine Tarrega, Rocío Coloma, Michela Falqui, Emma K. Luhmann, Lilliana Radoshevich, Susana Guerra

**Affiliations:** Department of Preventive Medicine, Public Health and Microbiology, Universidad Autónoma de Madrid, E-28029 Madrid, Spain; Department of Microbiology and Immunology, University of Iowa Carver College of Medicine, Iowa City, IA, 52242, USA

**Author notes:** Corresponding author. Mailing address: Department of Preventive Medicine and Public Health, Universidad Autónoma, E-28029 Madrid, Spain. Phone (+34) 91/ 497-5440. Fax: (+34) 91/ 497-5353. These authors contributed equally to this work and share first authorship (Martina Bécares, Manuel Albert).

## Abstract

Viruses have developed many different strategies to counteract immune responses, and *Vaccinia virus* (VACV) is one of a kind in this aspect. To ensure an efficient infection, VACV undergoes a complex morphogenetic process resulting in the production of two types of infective virions: intracellular mature virus (MV) and extracellular enveloped virus (EV), whose spread depends on different dissemination mechanisms. MVs disseminate after cell lysis, whereas EVs are released or propelled in actin tails from living cells. Here we show that ISG15 participates in the control of VACV dissemination. Infection of *Isg15-/-* mouse embryonic fibroblasts with VACV International Health Department-J (IHD-J) strain resulted in decreased EV production, concomitant with reduced induction of actin tails and the abolition of comet-shaped plaque formation, comparing with *Isg15+/+* cells. Transmission electron microscopy revealed accumulation of intracellular and a decrease in extracellular virus particles in the absence of Interferon Stimulated Gene 15 (ISG15), consistent with altered virus egress. Immunoblot and quantitative proteomic analysis of sucrose gradient-purified virions from both genotypes reported differences in protein levels and composition of viral proteins present on virions, suggesting an ISG15-mediated control of viral proteome. Last, the generation of a recombinant IHD-J expressing V5-tagged ISG15 (IHD-J-ISG15) allowed us to identify several viral proteins as potential ISG15 targets, highlighting the proteins A34 and A36, essential for EV formation. Altogether, our results indicate that ISG15 is an important host factor in the regulation of VACV dissemination.

**Author Summary:** Viral infections are a constant battle between the virus and the host. While the host’s only goal is victory, the main purpose of the virus is to spread and conquer new territories at the expense of the host’s resources. Along millions of years of incessant encounters, Poxviruses have developed a unique strategy consisting in the production two specialized “troops”: intracellular mature virions (MVs) and extracellular virions (EVs). MVs mediate transmission between hosts, and EVs ensure advance on the battlefield mediating the long-range dissemination.

The mechanism by which the virus ‘decides’ to shed from the primary site of infection and its significant impact in viral transmission is not yet fully established. Here, we demonstrate that this process is finely regulated by ISG15/ISGylation, an interferon-induced ubiquitin-like protein with broad antiviral activity. Studying the mechanism that viruses use during infection could result in new ways of understanding our perpetual war against disease and how we might win the next great battle.

## Introduction

Viruses are responsible for many infectious diseases, from mild illnesses such as the common cold, flu, and warts, to severe diseases such as acquired immunodeficiency syndrome (AIDS), Ebola hemorrhagic fever and the novel *Coronavirus* disease-19 (COVID-19). The current severe acute respiratory syndrome-*Coronavirus*-2 (*SARS-CoV-2*) pandemic is a good example of how viruses can cause global outbreaks with high mortality rates. In this sense, understanding the molecular mechanisms that operate during viral infections is essential for the development of efficient antiviral therapies against emerging viruses.

The *Interferon* (IFN)*-Stimulated Gene 15* (*Isg15*) encodes a small ubiquitin-like post-translational modifier that regulates a plethora of cellular pathways through the modulation of the proteome. ISG15 exerts its functions by covalent conjugation to target proteins in a process termed ISGylation (1), or as a free molecule both inside and outside the cell (2). Intracellular unconjugated ISG15 controls the stability of target proteins through non-covalent interactions (3-5), whereas extracellular ISG15 acts as a cytokine and modulates immune cell functions by binding to lymphocyte function-associated antigen 1 (LFA-1) (6-8). ISG15 has a well-established antiviral function against diverse biomedically relevant viruses, including *Human Immunodeficiency virus* (HIV), *Influenza virus, SARS-CoV-2*, and *Human Herpesvirus (*HSV-1) (9-12). Such antiviral activity is achieved through direct interaction of ISG15 with viral proteins, or through the modulation of host proteins to limit the progression of the infection and to enhance immune responses (9). Nevertheless, viruses have developed different strategies to counteract the action of ISG15, including the cleavage of ISG15 from ISGylated proteins (11, 13), the sequestration of ISGylated viral proteins (14), or the blockage of ISGylation of target proteins (15, 16).

The *Poxviridae* is a family of large, enveloped, linear double-stranded DNA viruses that replicate entirely in the cytoplasm of infected cells. A unique feature of the *Poxviridae* family is the production of two distinct infectious forms: intracellular mature virus (MV) and extracellular enveloped virus (EV) (17, 18). MVs mediate host-to-host transmission, whereas EVs disseminate within the host causing systemic infection (19). EVs derive from MVs that bud from the plasma membrane (20), or that undergo wrapping by the trans-Golgi network (TGN) or endosomal membranes (21, 22). In this process, an intermediate virus form, the intracellular enveloped virus (IEV), is generated. IEVs are transported on microtubules towards the plasma membrane, which they fuse with to release EVs to the extracellular medium (23, 24); however, a variable percentage of EVs remains attached to the plasma membrane as cell-associated enveloped virus (CEV). This membrane fusion event causes IEV outer membrane-associated proteins to remain in the plasma membrane, where they induce the polymerization of actin tails to promote virion spread to neighboring cells (18). EVs are not just MV particles enclosed within a second lipid membrane, as they differ in protein composition. In *Vaccinia virus* (VACV), the proteins A25 and A26 are exclusive of MVs, whereas the proteins A36, F12, E2, B5, A34, F13, A56 and K2 are exclusive of wrapped forms (IEV and EV) (25). The production of each virus form varies among the different strains of VACV. For example, in infections with VACV Western Reserve strain (WR), MVs represent the majority of infectious progeny, whereas VACV International Health Department-J strain (IHD-J) produces a high EV/MV ratio as a result of a point mutation (K151Q) in the A34R gene, involved in EV formation and dissemination (26).

Previous research demonstrated that ISG15 has a role in the control of VACV infection (27, 28), and that VACV E3 protein counteracts the action of ISG15 (15, 27). Moreover, we previously showed that VACV reduces mitochondrial respiration of macrophages in an ISG15-dependent manner (29). Several lines of evidence argue that VACV uses an exosome-like pathway for EV formation and release (25, 30). Considering that ISG15 has been shown to impair exosome secretion by promoting the fusion of multivesicular bodies with lysosomes (31), we sought to explore whether the ISG15/ISGylation system has an impact in VACV EV formation and dissemination. Here, we show that ISG15 modulates VACV dissemination and that its absence alters the virion proteome. Infection of *Isg15-/-* mouse embryonic fibroblasts (MEF) with VACV IHD-J resulted in reduced EV production, consistent with reduced actin tail formation, the accumulation of virus particles in the cytoplasm, and the abolition of comet-shaped plaques, compared with *Isg15+/+* MEF. In addition, a quantitative proteomic analysis of purified virions from *Isg15-/-* MEF reported and enrichment in proteins of both MVs and wrapped virions, confirming the accumulation of distinct virus forms in *Isg15-/-* MEF. Last, the generation of a recombinant virus expressing V5-tagged ISG15 (IHD-J-ISG15) allowed us to identify viral proteins that potentially interact with ISG15, highlighting the protein A36, essential for the formation of actin tails (18).

In summary, our study contributes to the comprehension of how the ISG15/ISGylation system modulates VACV infection, and reinforces the idea of ISG15 as an essential host factor in the coordination of viral pathogenesis and antiviral immune responses.

## Results

### ISG15 is required for EV dissemination during IHD-J infection

To analyze whether ISG15 has a role in EV release, we evaluated the formation of comet-like plaques during IHD-J infection. *In vitro*, EVs disseminate by convection forming comet-shaped plaques, consisting of a central lysis plaque (the comet ‘head’) and an ensemble of secondary satellite plaques (the comet ‘tail’) (32), what allows to easily evaluate an enhanced production of EVs. Monolayers of *Isg15+/+* and *Isg15-/-* MEF were infected with IHD-J (0.0001 plaque forming units (PFU)/cell) and incubated in liquid medium. At 48 hours post-infection (hpi), monolayers were fixed and stained with 0.2% crystal violet in 10% formaldehyde to visualize lysis plaques. Surprisingly, a drastic reduction in the formation of comet-shaped plaques was observed in *Isg15-/-* MEF (Fig. 1A), indicating that EV release is impaired in the absence of ISG15.

**Figure 1.**
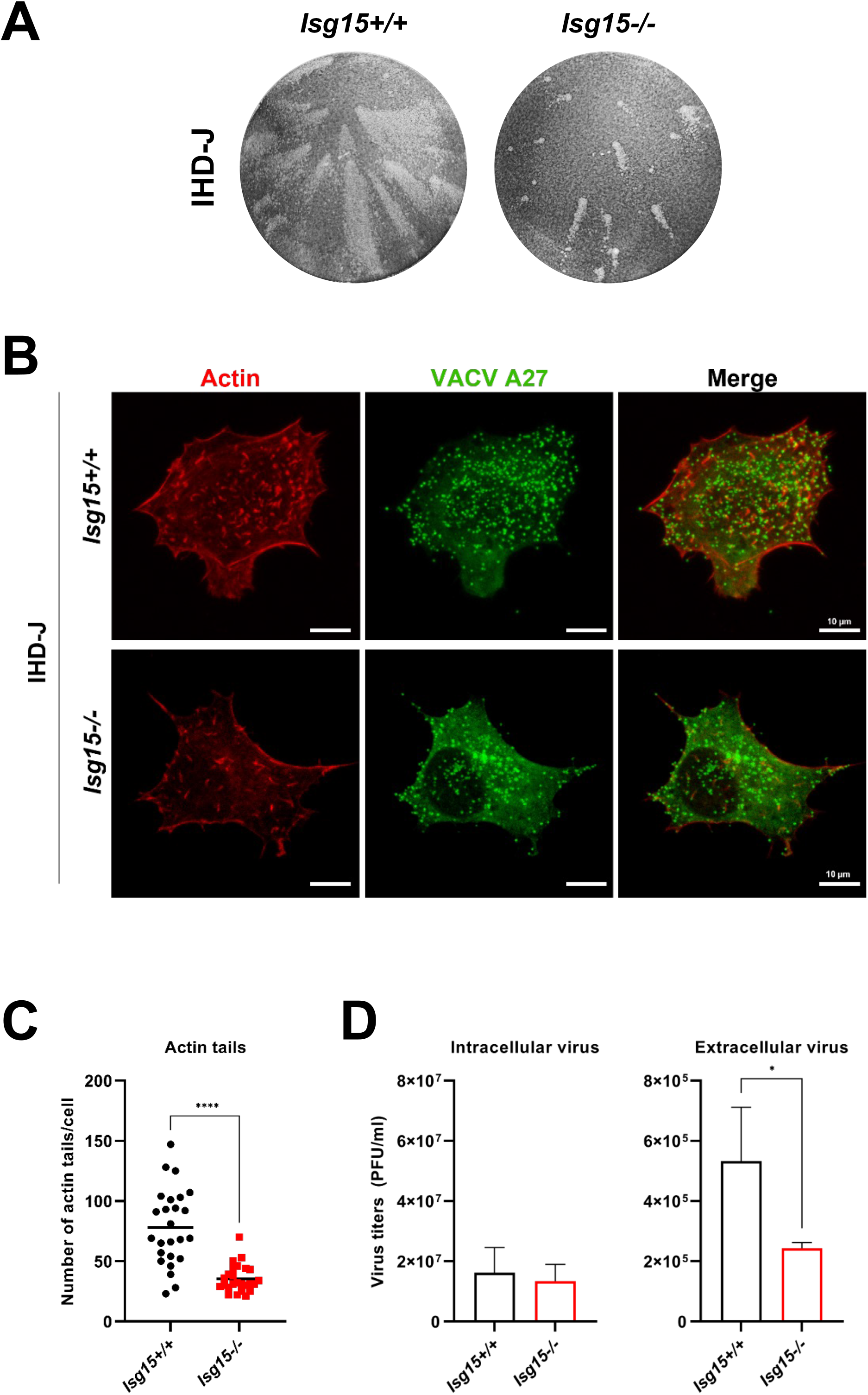
The absence of ISG15 impairs comet-shaped plaque, actin tail and EV production. **(A) Comet-shaped plaque formation is dramatically reduced in *ISG15-/-*MEFs**. Monolayers of immortalized *ISG15+/+* and *ISG15-/-*MEFs were infected with IHD-J (0.0001 PFU/cell). At 48 hpi, cells were fixed and stained with 0.2% crystal violet in 4% PFA for comet-shaped plaque visualization **(B, C) Actin tail formation is significantly reduced in *ISG15-/-* MEFs**. Immortalized *ISG15+/+* and *ISG15-/-*MEFs growing in coverslip were infected with IHD-J (2 PFU/cell). At 9 hpi, actin tails were visualized by phalloidin-staining (red), and viral particles were visualized using an anti-A27 antibody, followed by a fluorescent secondary antibody (green). Actin tails were quantified using Leica AF software (N = 25, each genotype), and actin tail numbers (Mean ± SD) of individual cells are represented. **(D) EV release is significantly decreased in *ISG15-/-*MEFs**. Immortalized *ISG15+/+* and *ISG15-/-*MEFs were infected with IHD-J (2 PFU/cell). At 16 hpi, infectious viral particles from supernatants (extracellular virus) and cell extracts (intracellular virus) were titrated by plaque assay. Mean ± SD values from three independent experiments are represented. *, *P* value < 0.05; **, *P* value < 0.01; ***, *P* value < 0.005; **** *P* value < 0.0001.

To study whether the reduction in comet-shaped plaques was cell-line specific, we silenced *Isg15* in NIH/3T3 cells using two specific lentiviral vectors (L1 and L2) expressing *Isg15* shRNAs. A clear reduction of *Isg15* mRNA was observed in type I IFN-treated cells transduced with both shRNAs in comparison with non-transduced cells, and with cells transduced with a control shRNA (scramble; SC) (Supp. Fig.1A). Western blot analysis further confirmed the interference of *Isg15* expression, as ISG15 was undetectable in L1- and L2-transduced cells (Supp. Fig.1B), whereas a clear induction of the protein was observed in IFN-treated control cells. *Isg15* interference in L1- and L2-transduced NIH/3T3 cells abolished the formation of comet-shaped plaques, validating our results obtained in immortalized MEF (Supp. Fig.1C).

VACV relies on the cytoskeleton for dissemination. Wrapped virions are transported on microtubules to the cell surface, and CEVs induce the polymerization of actin and the formation of actin tails that propel virions to neighboring cells (17). ISG15 has been shown to modulate actin cytoskeleton dynamics (33), and several cytoskeleton proteins were identified as ISGylation target (10). Therefore, we explored whether the absence of ISG15 alters actin tail formation during IHD-J infection. *Isg15+/+* and *Isg15-/-* MEF were infected (2 PFU/cell, 9 h) and samples were processed for immunofluorescence analysis. Actin and MVs were labeled using Alexa Fluor™ 594-phalloidin and anti-VACV A27 protein-specific antibodies, respectively, and cells were imaged by confocal microscopy. Actin tails were detected in both *Isg15+/+* and in *Isg15-/-* MEF, although a significant reduction in the number of actin tails was observed in *Isg15-/-* MEF, indicating a role of ISG15/ISGylation in the modulation of actin tail formation (Fig. 1B and 1C). The reduction in the formation of actin tails and comet-shaped plaques was consistent with reduced extracellular virus titers in infected *Isg15-/-* MEF (2 PFU/cell, 16 h) (Fig. 1D, right panel). Titers of intracellular virus, however, did not significantly differ between genotypes (Fig. 1D, left panel).

To rule out a delay in the infection of *Isg15-/-* MEF as the cause of reduced actin tail formation and EV release, we analyzed the levels of early (E3) and intermediate (A27 and F13) viral proteins by western blot in infected *Isg15+/+* and *Isg15-/-* MEF (2 PFU/cell) at 9 and 16 hpi. As well, we analyzed the subcellular distribution of A27 and F13 proteins by confocal microscopy at 9 hpi. Neither the levels (Fig. 2A) nor the distribution (Fig. 2B) of any of the viral proteins analyzed significantly differed between *Isg15-/-* and *Isg15+/+* MEF, indicating that the infection occurred normally in both genotypes using the cascades mechanism as described (34)

**Figure 2.**
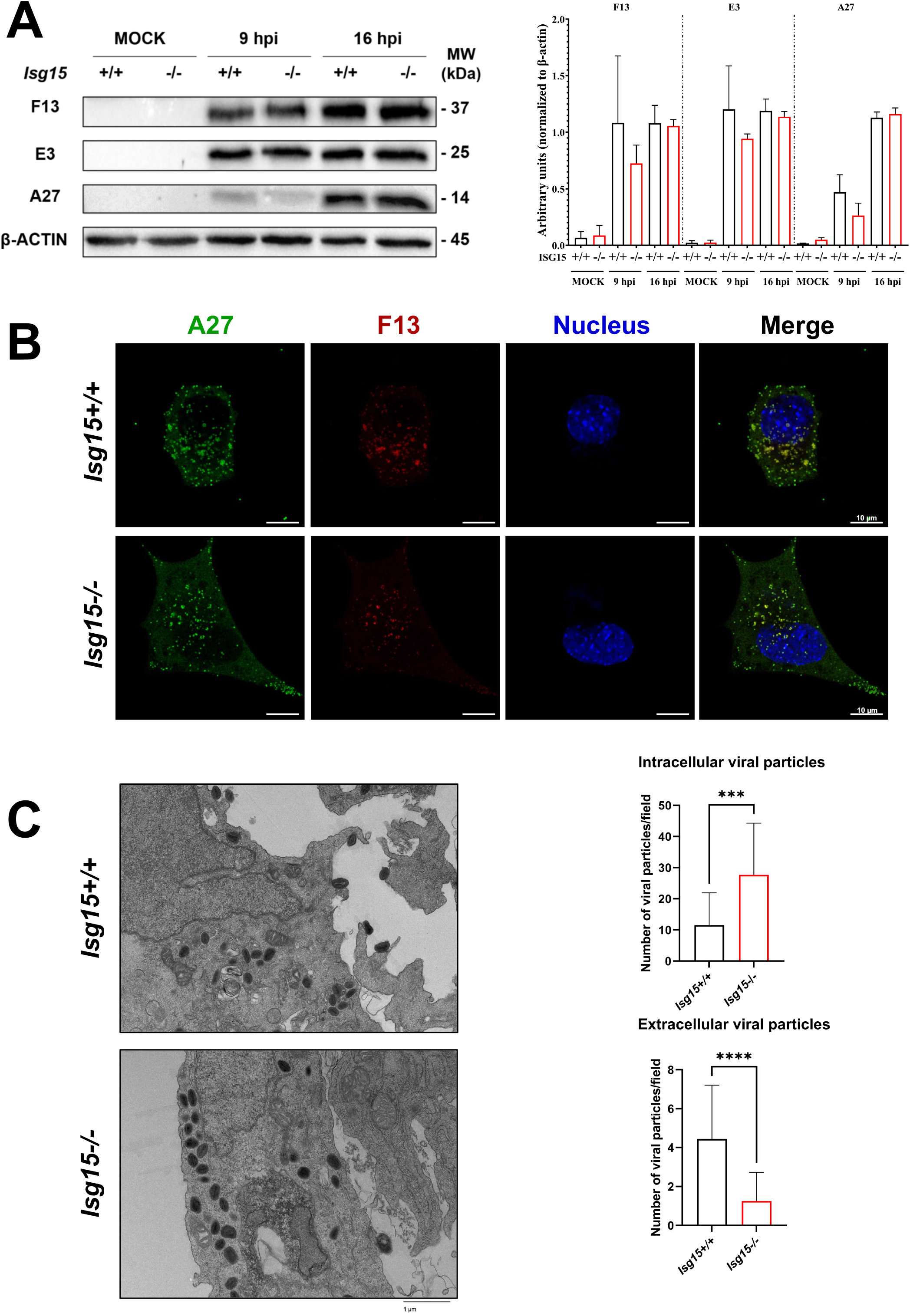
IHD-J morphogenetic process is altered in *ISG15-/-*MEFs. **(A) Viral protein synthesis is not affected by the absence of ISG15**. Immortalized *ISG15+/+* and *ISG15-/-*MEFs were infected with IHD-J (2 PFU/cell) and, at the indicated times post-infection, equal amounts of proteins from cell extracts were analyzed by Western Blot. Specific antibodies for VACV early E3 (*E3L*) and late proteins F13 (*F13L*), A27 (*A27L*) were used. Actin was used as loading control. Molecular weights (MW) in kilodaltons (kDa) are indicated, based on protein standards. Protein levels of 3 independent experiments were analyzed with Fiji software. Mean ± SD are indicated. **(B) Viral protein distribution does not change between *ISG15+/+* and *ISG15-/-* MEFs**. Confocal microscopy analysis of immortalized *ISG15+/+* or *ISG15-/-*MEFs infected with IHD-J (2 PFU/cell) at 9 hours post-infection. Viral proteins A27 (green) and F13 (red) were labeled by immunofluorescence; DAPI (4′,6-diamidino-2-phenylindole) (blue) was used to stain the nuclear DNA. **(C, D) The absence of ISG15 causes accumulation of intracellular virions and a diminution in extracellular virus**. Representative images of *ISG15+/+* and *ISG15-/-* MEFs infected with IHD-J (2 PFU/cell; 9 h) are shown. Intracellular or extracellular viral particles per field were quantified with Fiji software (2500x magnification, N = 20, each genotype). Mean ± SD are indicated. *, *P* value < 0.05; **, *P* value < 0.01; ***, *P* value < 0.005; **** *P* value < 0.0001.

In addition, we analyzed intracellular virus particles by transmission electron microscopy (TEM) of infected MEF (2 PFU/cell, 9 h) to assess whether *Isg15-/-* MEF accumulated more virus particles in the cytoplasm due to defective virus release. Interestingly, this approach revealed increased number of intracellular virus particles and a clear decrease of extracellular virus was observed of *Isg15-/-* MEF (Fig. 2C and 2D). These observations were consistent with defects in EV production and dissemination in *Isg15-/-* MEF and suggested that some of these intracellular viral particles might be defective, considering that the titers of intracellular infectious virus did not significantly differ between genotypes (Fig. 1D, left panel).

Last, to determine whether the effect of ISG15 on VACV dissemination was exclusive of IHD-J or it also affected other VACV strains, we extended our study to the WR strain. We evaluated the formation of comet-shaped plaques in WR-infected MEF (0.0001 PFU/cell, 72 h), and observed a reduction in comet-shaped plaques in *Isg15-/-* MEF (Supp. Fig. 2A). As well, the formation of actin tails was also impaired in these cells (Supp. Fig. 2B), indicating that the role of ISG15 in VACV dissemination is not strain-specific.

Altogether, our results point out the relevance of ISG15 in VACV dissemination, demonstrating that the ISG15/ISGylation system is involved in the regulation of virion egress, modulating the formation actin tails and EV release.

### Protein composition of VACV progeny is altered in the absence of ISG15

To determine whether the alterations in VACV dissemination could be caused by changes in virion proteins due to the absence of ISG15, we studied how ISG15 modulates the proteome of VACV. For that purpose, we performed a quantitative proteomic analysis with sucrose gradient-purified virions obtained from infected *Isg15-/-* and *Isg15+/+* MEF. This analysis allowed us to identify and quantify numerous viral proteins whose expression could be modulated by ISG15, present in purified virions. Overall, we identified 717 proteins, of which 402 were reliably identified (present in all 3 biological replicates; False Discovery Rate (FDR) < 0.05). Of these 402 proteins, only 177 proteins were reported as significantly upregulated or downregulated (FDR < 0.01) in virions from *Isg15-/-* MEF, comparing with those from *Isg15+/+* MEF. We found quantitative differences in the levels of 63 viral proteins comparing virions from both genotypes (Table 1). Interestingly, 58 viral proteins were significantly enriched in virions purified from *Isg15-/-* MEF, whereas 5 (N1, A34, F11, D13 and IL8B) were significantly reduced (Fig. 3A).

**Table 1.**
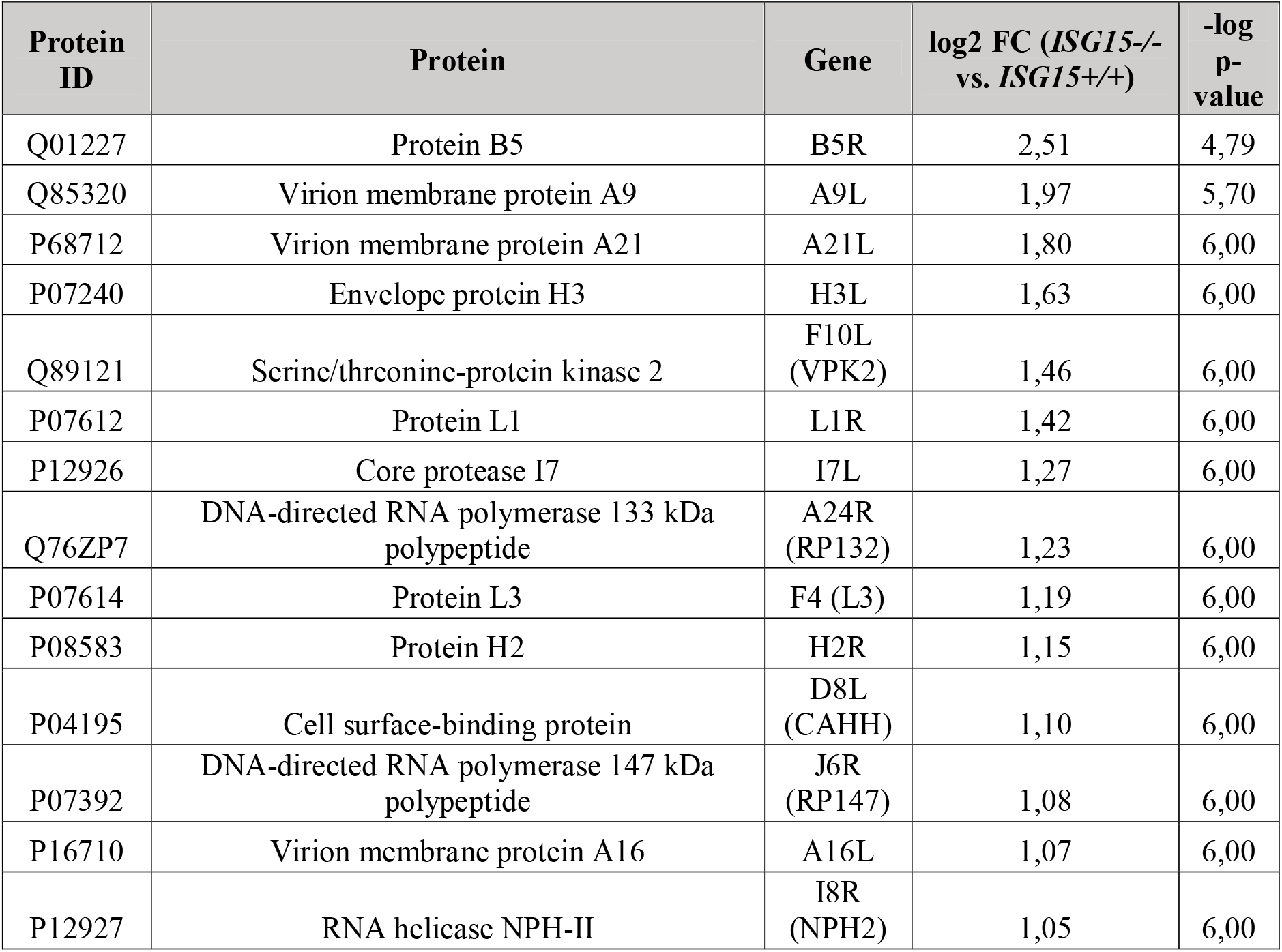
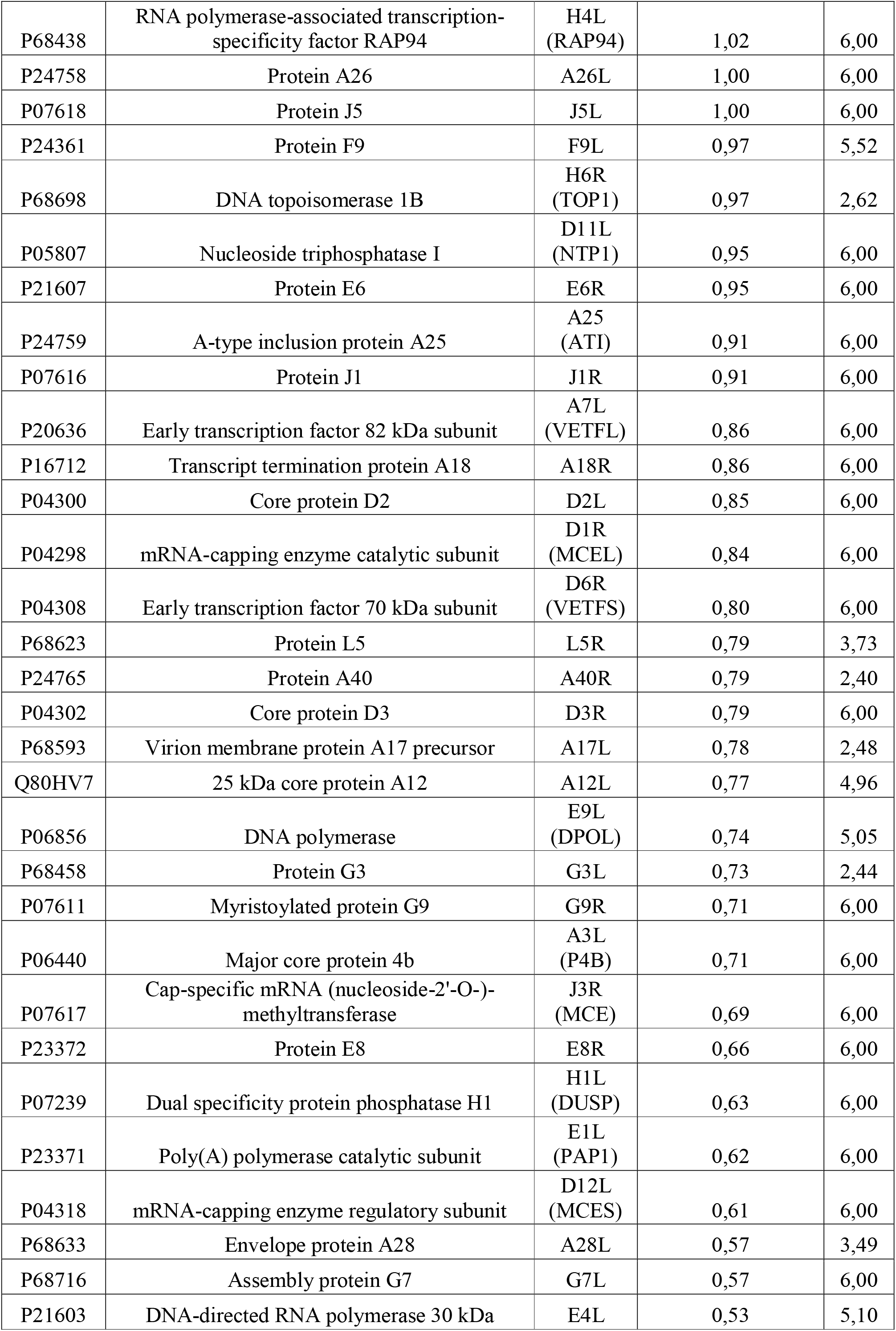
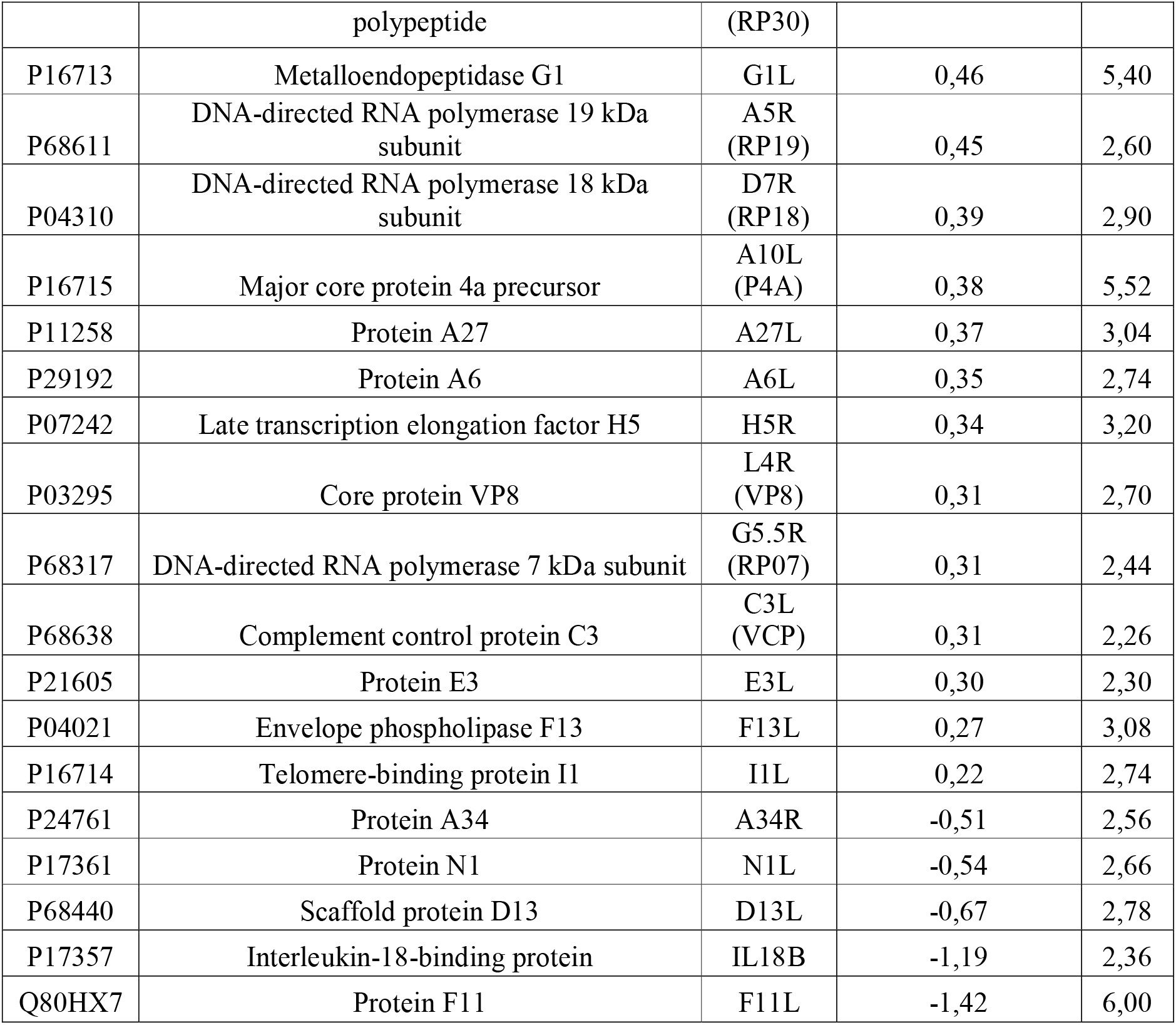
Viral protein signature of highly purified virions from *ISG15+/+* and *ISG15-/-* MEFs. Comparison of viral proteins present in virions purified from *ISG15-/-*vs. *ISG15+/+* MEF, identified and quantified by LC-ESI-MS/MS. To identify proteins significantly enriched in each sample, a t-test was performed (FDR = 0.05 and S_0_ =1). Table 1 lists: Protein ID (column A); protein name (column B); gene name (column C); log2 fold change (FC) (*ISG15-/-vs. ISG15+/+*) of the levels of each protein (column D), and statistical significance (−log P-value) (column E). Proteins are ordered from most enriched (top) to less enriched (bottom) in *ISG15-/-* samples.

**Figure 3.**
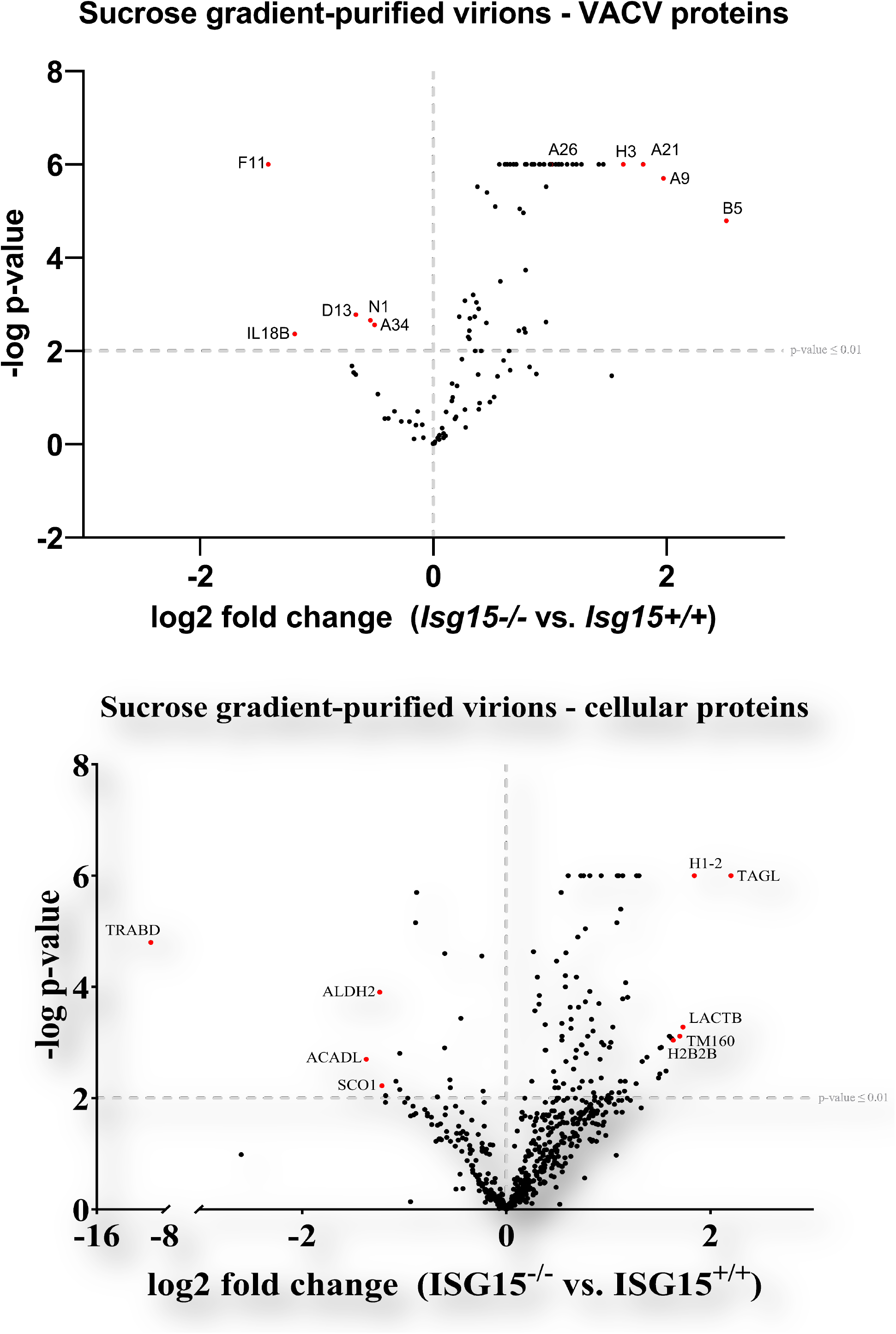
The proteomes of purified virions change in the absence of ISG15. **(A) The viral protein content of highly purified virions from *ISG15+/+* and *ISG15-/-*MEF differs between genotypes**. Immortalized *ISG15+/+* or *ISG15-/-*MEFs were infected with IHD-J (0.01 PFU/cell, 48 h), and intracellular virions were purified by ultracentrifugation through a 20% sucrose cushion followed by ultracentrifugation through a sucrose gradient (20-45%). Purified virions were processed for LC-MS/MS analysis. Viral proteins are represented in the volcano plot. A *P* value ≤ 0.01 cut-off is indicated in the graph. Proteins of interest significantly enriched in virions isolated from *ISG15+/+* cells (upper left) or *ISG15-/-*cells (upper right) are labeled and highlighted in red. **(B) The cellular protein content of highly purified virions from *ISG15+/+* and *ISG15-/-*MEF differs between genotypes**. Immortalized *ISG15+/+* or *ISG15-/-*MEFs were infected with IHD-J (0.01 PFU/cell, 48 h), and intracellular virions were purified by ultracentrifugation through a 20% sucrose cushion followed by ultracentrifugation through a sucrose gradient (20-45%). Purified virions were processed for LC-MS/MS analysis. Cellular proteins are represented in the volcano plot. A *P* value ≤ 0.01 cut-off is indicated in the graph. The most significantly enriched proteins in virions isolated from *ISG15+/+* cells (upper left) or *ISG15-/-*cells (upper right) are labeled and highlighted in red.

Among the enriched proteins in virions from *Isg15-/-* MEF we found proteins specific of wrapped virus forms, such as B5 and F13, and the protein A26, specific of MVs. These observations were consistent with defects in EV formation and the accumulation of diverse virus forms in *Isg15-/-* MEF. As well, we detected a significant reduction in the levels of A34 and F11 in virions from *Isg15-/-* MEF. A34 is involved in actin tail formation, EV release, the disruption of EV membranes prior to virus entry, and the localization of the proteins B5, A33 and A36 to wrapped virions (35). As well, the protein F11 promotes viral spread by modulating actin cytoskeleton through interactions with myosin 9A and RhoA (36, 37). Thus, reduced levels of A34 and F11 in virions from *Isg15-/-* MEF might be related to the defects in VACV dissemination in these cells.

We also observed quantitative differences in cellular proteins fingerprint detected in the protein extracts co-migrating with the purified virions from infected *Isg15-/-* and *Isg15+/+* MEFs. Specifically, 108 cellular proteins were reliably identified and significantly differed between both types of virions (FDR < 0.01). Of the total, 15 host proteins showed lower levels in virions purified from *Isg15-/-* MEF, while the rest were significantly upregulated (Table 2 and Fig. 3B). Interestingly, most of the proteins identified were related to mitochondrial metabolism and, particularly, to oxidative phosphorylation, what might be explained by increased synthesis of mitochondrial proteins to support the large consumption of ATP required for viral production and dissemination (Supp. Fig. 3).

**Table 2.**
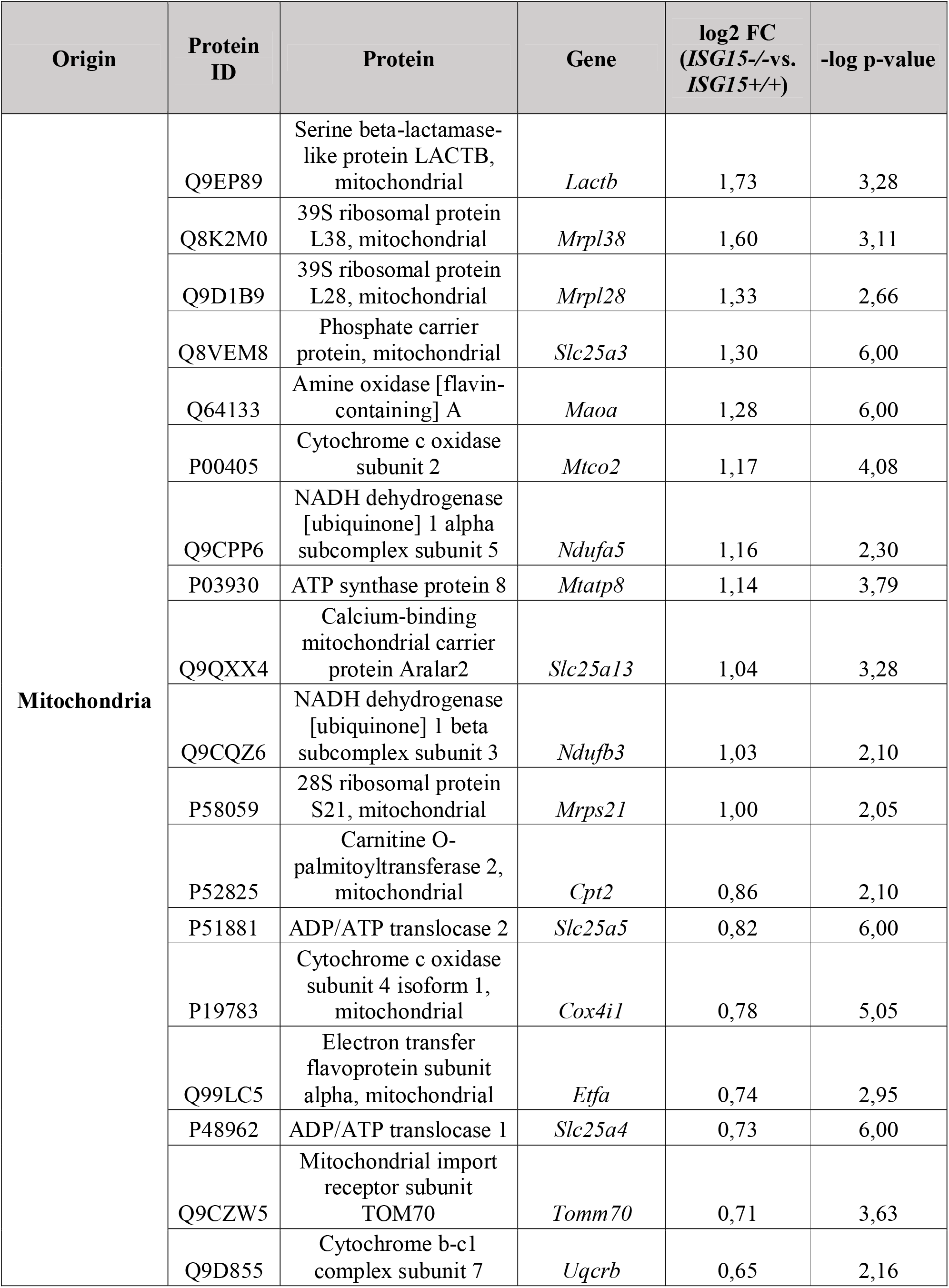
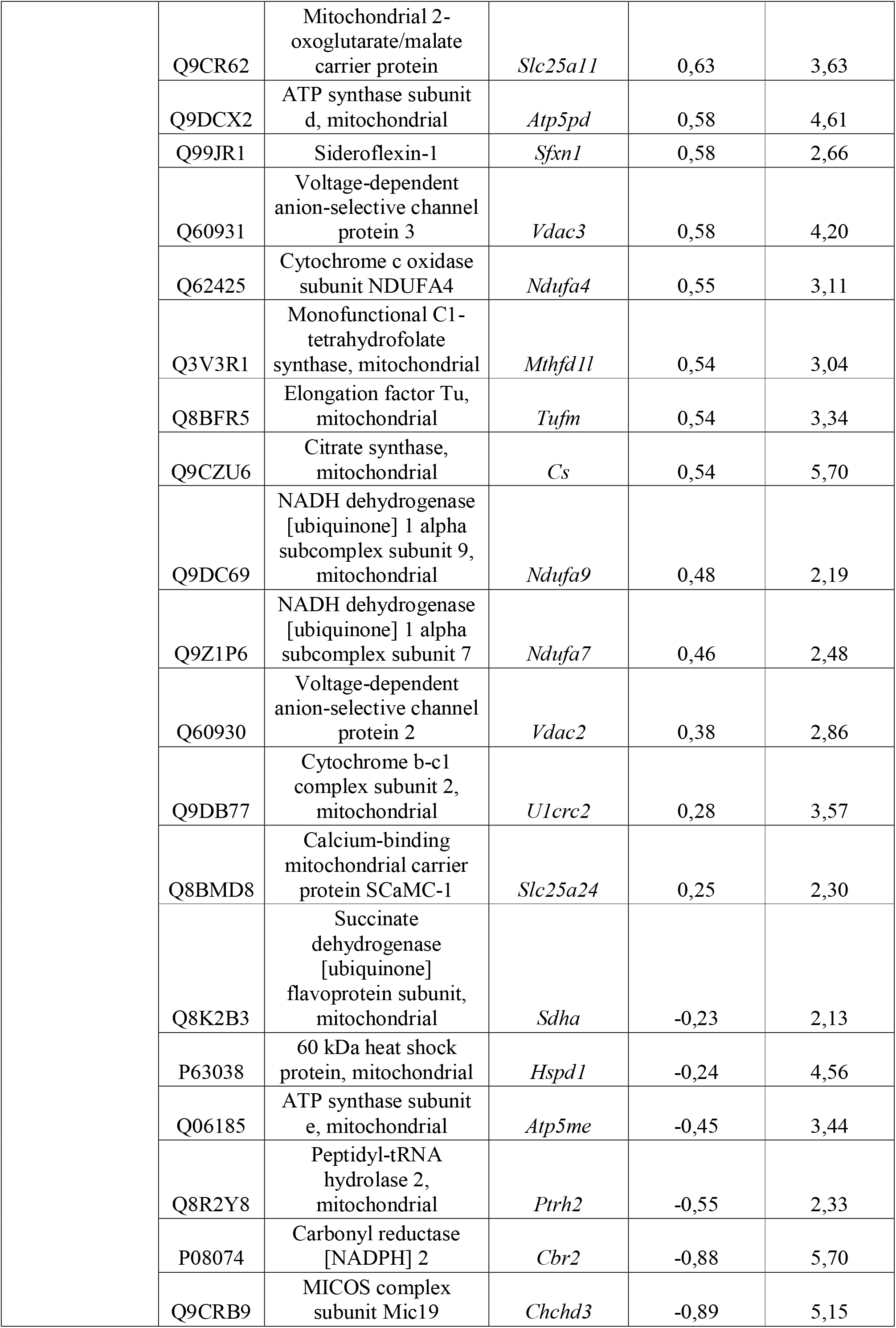
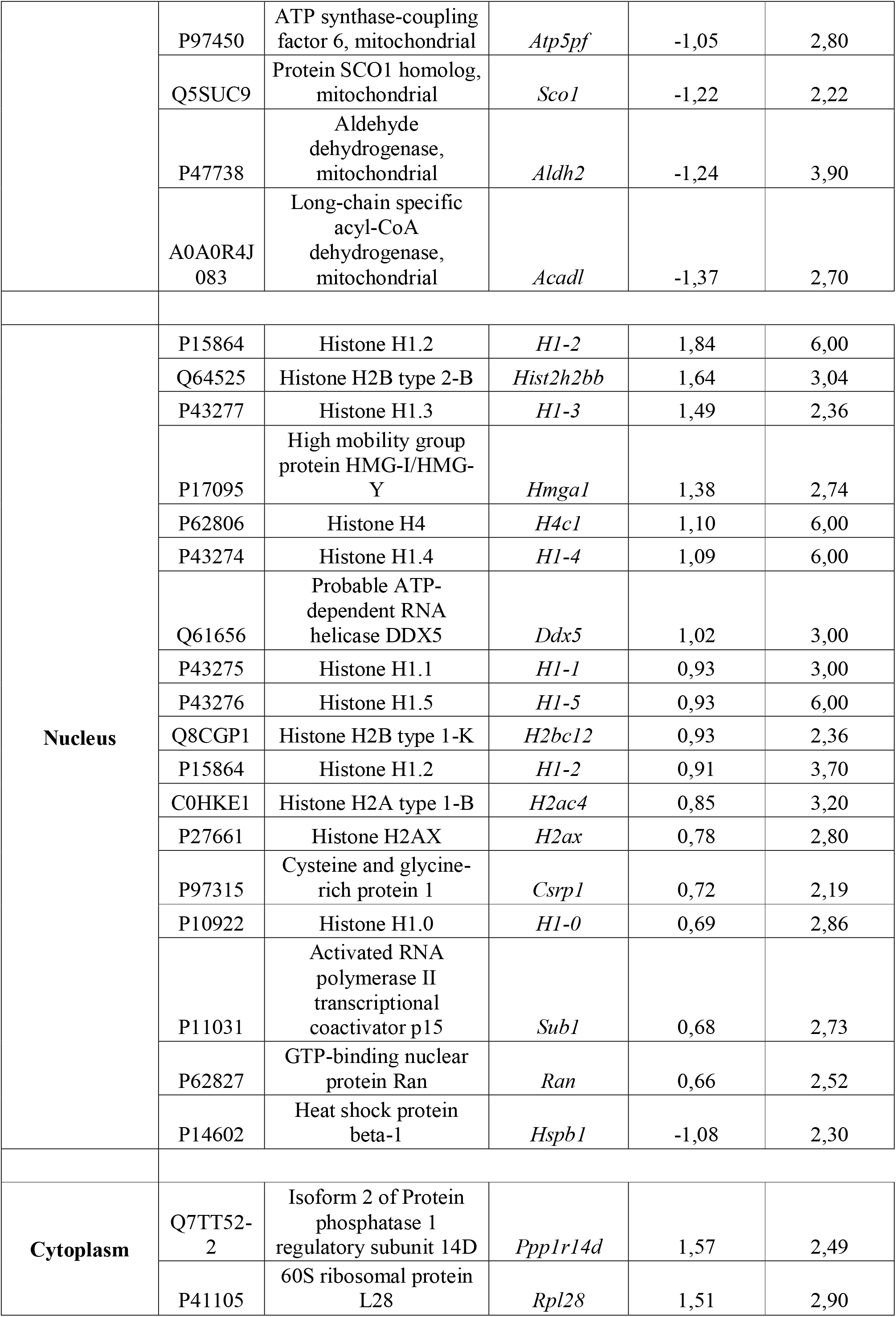
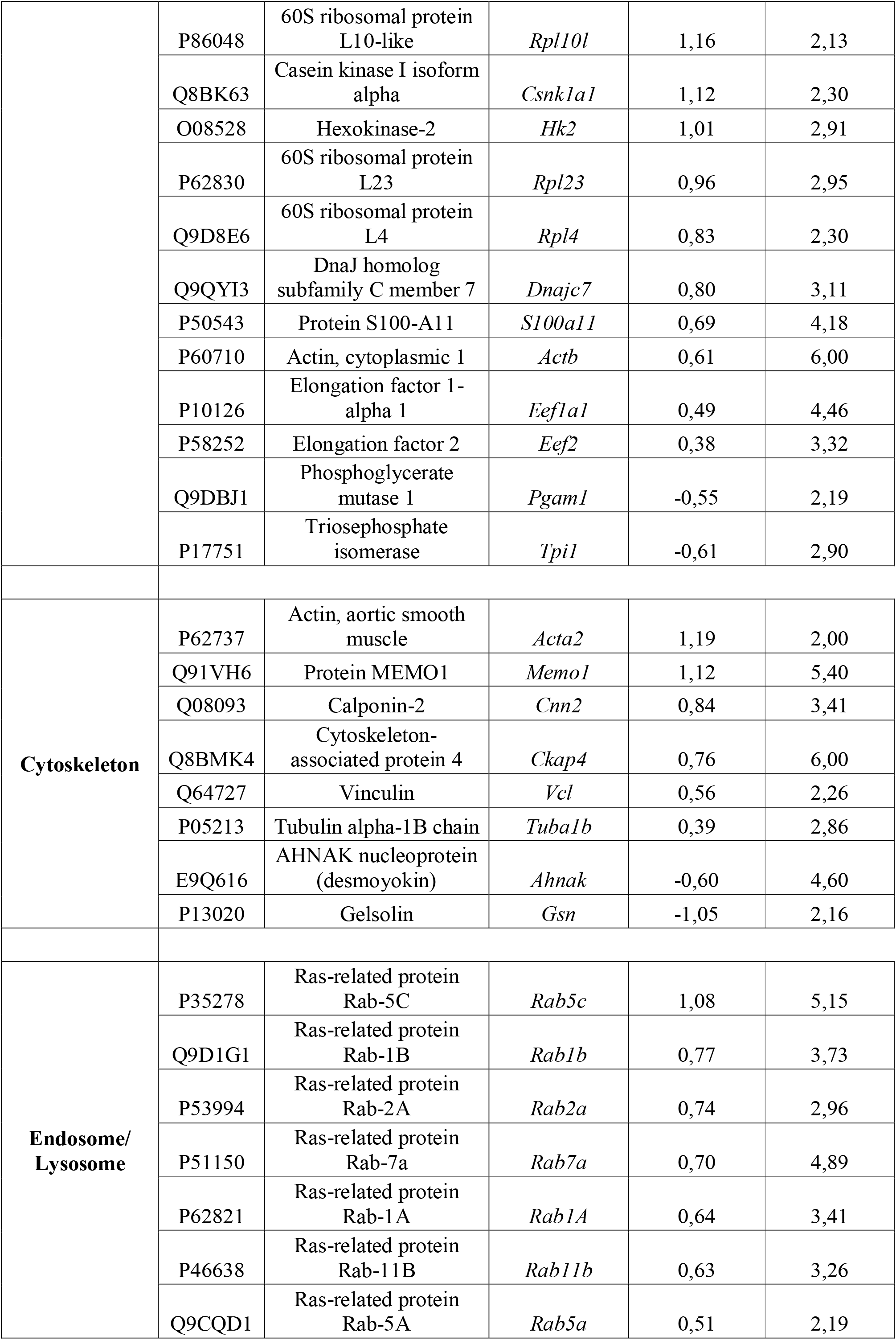
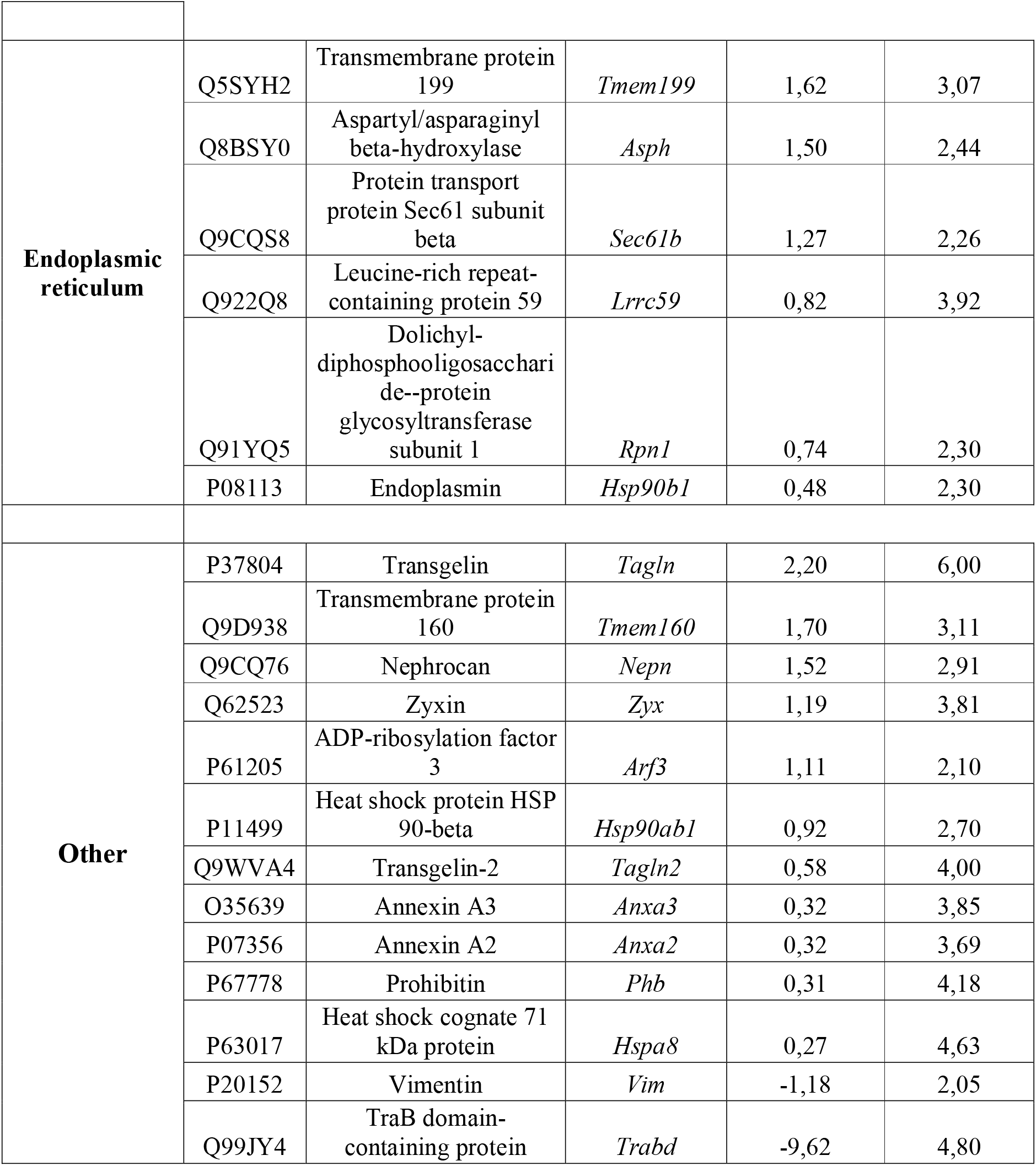
Cellular protein signature of highly purified virions from *ISG15+/+* and *ISG15-/-* MEFs. Comparison of cellular proteins present in virions purified from *ISG15-/-vs. ISG15+/+* MEF, identified and quantified by LC-ESI-MS/MS. To identify proteins significantly enriched in each sample, a t-test was performed (FDR = 0.05 and S_0_ =1). Table 2 lists: Protein subcellular location (referred as “Origin”) (column A); protein ID (column B); protein name (column C); gene name (column D); log2 fold change (FC) (*ISG15-/-vs. ISG15+/+*) of the levels of each protein (column E), and statistical significance (−log P-value) (column F). Proteins are ordered from most enriched (top) to less enriched (bottom) in *ISG15-/-*samples within each “Protein origin” group.

### ISG15 binds VACV proteins

In our mission to determine how ISG15 modulates VACV proteome, we generated a recombinant virus expressing murine ISG15 (IHD-J-ISG15) to assess whether ectopically expressed ISG15 binds VACV proteins when co-expressed. To distinguish endogenous ISG15 from that expressed by the recombinant virus, the latter was tagged with a V5 epitope at its N-terminus. The V5-ISG15 construct was inserted into the VACV thymidine kinase (TK) locus of parental IHD-J, under the transcriptional control of the VACV Early/Late romoter. The IHD-J-ISG15 virus was generated following an infection-transfection protocol (38) and its genome is depicted in Fig. 4A. The correct insertion and purity of recombinant IHD-J-ISG15 viruses were analyzed by PCR using primers annealing in the VACV TK-flanking regions that confirmed the presence of the full-length murine ISG15 gene. Moreover, the correct sequence of full-length ISG15 gene inserted in the VACV TK locus was also confirmed by DNA sequencing (not shown).

**Figure 4.**
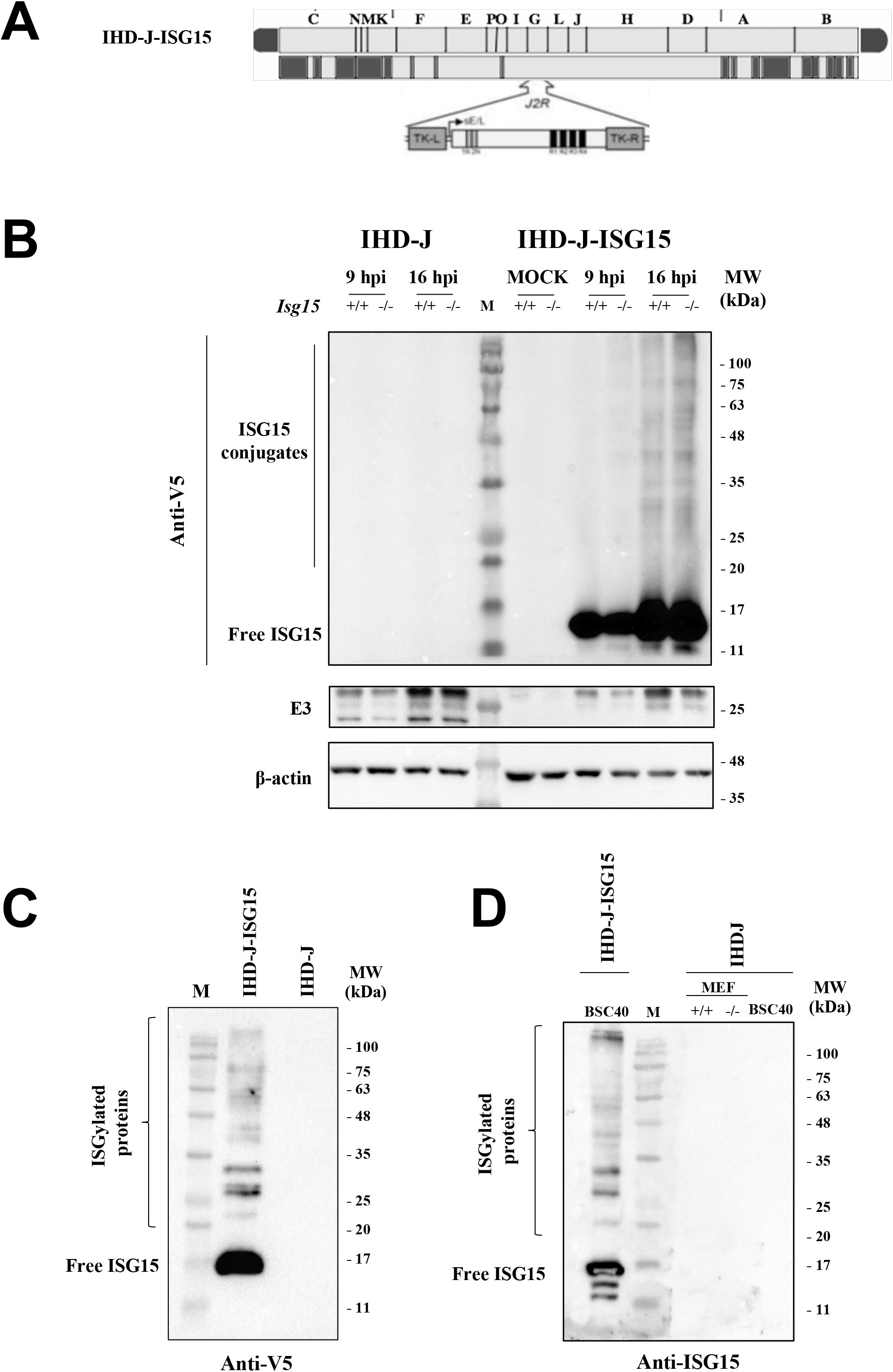
Generation and *in vitro* characterization of IHD-J-ISG15. **(A) Scheme of the IHD-J-ISG15 genome map**. The full-length murine ISG15 with a V5 epitope fused to its N-terminus was inserted within the VACV TK viral locus (*J2R*), and its expression was driven by the VACV E/L promoter **(B) Expression of V5-ISG15 fusion protein**. Immortalized *ISG15+/+* or *ISG15-/-* MEFs were mock infected or infected at 2 PFU/cell with IHD-J or IHD-J-ISG15. At the indicated times post-infection, proteins were separated by SDS-PAGE, and the expression of viral early (E3) and late (A27, A4) proteins was analyzed by Western Blot, using specific antibodies. Actin was used as loading control. **(C, D) Purified IHD-J-ISG15 virions contain ISGylated proteins and free ISG15**. Purified IHD-J or IHD-J-ISG15 from BSC40 cells or immortalized *ISG15+/+* and *ISG15-/-*MEF were fractionated by SDS-PAGE, and the expression of V5-ISG15 (C) and ISG15 (D) were analyzed by Western Blot, using specific antibodies. Molecular weights (MW) in kilodaltons (kDa) are indicated, based on protein standards (M).

To demonstrate that IHD-J-ISG15 constitutively expressed the full-length murine ISG15, *Isg15+/+* and *Isg15-/-*MEF were mock infected or infected with 0.5 PFU/cell of parental IHD-J or IHD-J-ISG15 for 9 and 16 h, and the expression of recombinant ISG15 was analyzed by western blot using an anti-V5 specific antibody. The results show that both monomeric ISG15 and ISGylated proteins were detected during IHD-J-ISG15 infection (Fig. 4B). Next, to determine whether virus-expressed ISG15 could be conjugated to virion proteins, sucrose gradient-purified IHD-J and IHDJ-ISG15 virions were analyzed by western blot. We detected protein ISGylation in protein extracts from purified intracellular virions, something we did not observe in IHD-J-infected *Isg15+/+* MEF, probably due to the low levels of ISG15 produced during VACV infection (Fig. 4C). Similar results were observed in MVs purified from BSC40 cells, *Isg15+/+* or *Isg15-/-* MEFs, using a specific ISG15 antibody (Fig. 4D).

To identify ISGylated viral proteins, we performed immunoprecipitation assays with IDH-J and IHD-J-ISG15 sucrose gradient-purified virions to capture and detect V5-ISG15 targets, using a V5 antibody. Western blot analysis of the immunoprecipitated extracts showed that several proteins reacted against an ISG15-specific antibody (Fig. 5A). To identify these proteins, the samples were subjected to Liquid Chromatography with Electrospray Ionization and Tandem Mass Spectrometry (LC-ESI-MS/MS). A high-resolution short gradient was performed, and 21 VACV proteins were identified in the immunoprecipitated extract (Table 3 and Fig. 5B). Several of the identified viral proteins are involved in the regulation of cytoskeleton dynamics (F11, A34, A36), virion morphogenesis (A6, A15, A19), and virion entry (A21). We further validated these results with the detection of A36 in the immunoprecipitated extracts from IHD-J-ISG15 virions (Fig. 5C). A36 is involved in actin tail formation (39), therefore, alterations in its localization and/or function due to the absence of ISG15 might explain the reduced actin tail formation in *Isg15-/-* MEF.

**Table 3.**
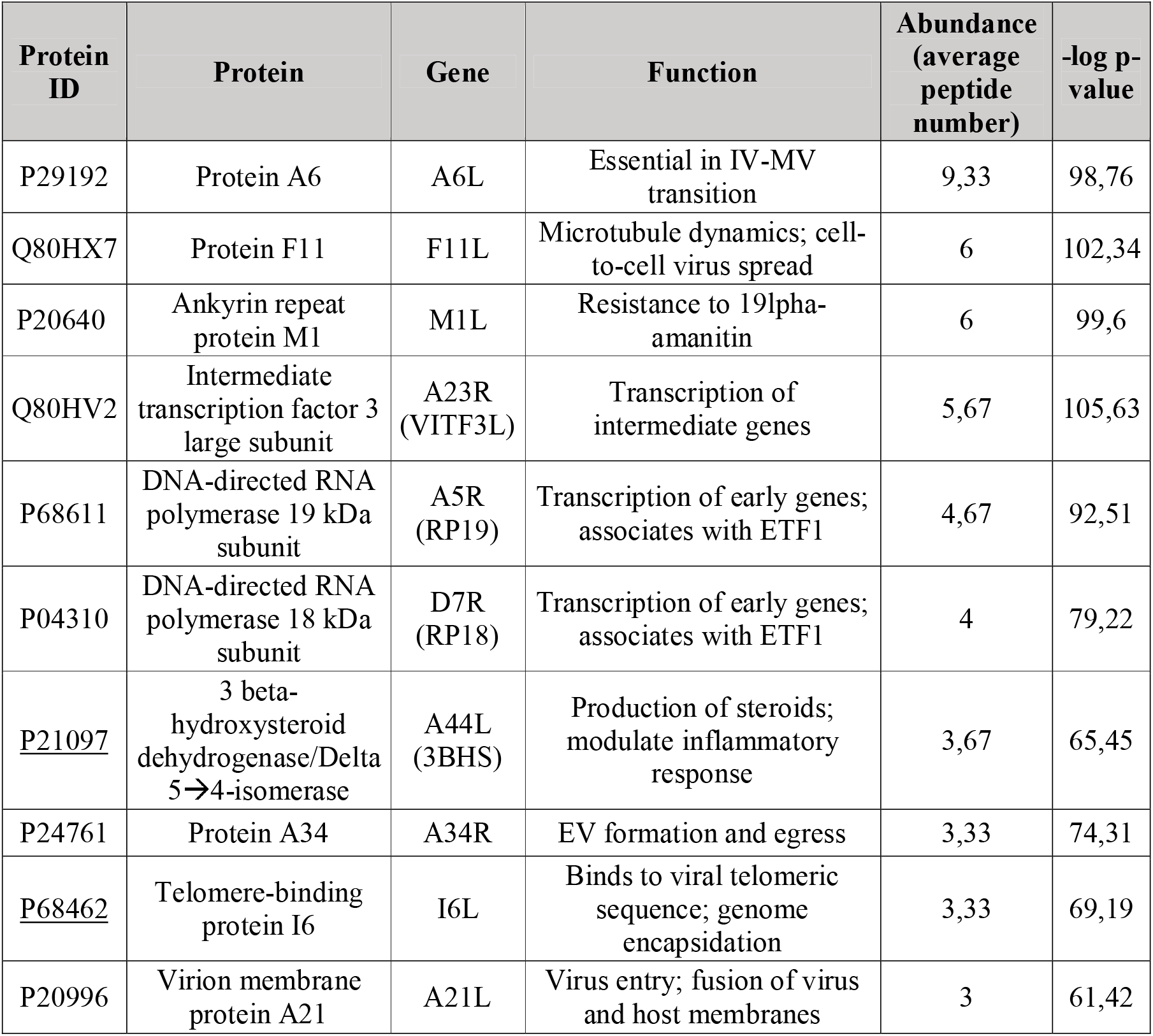
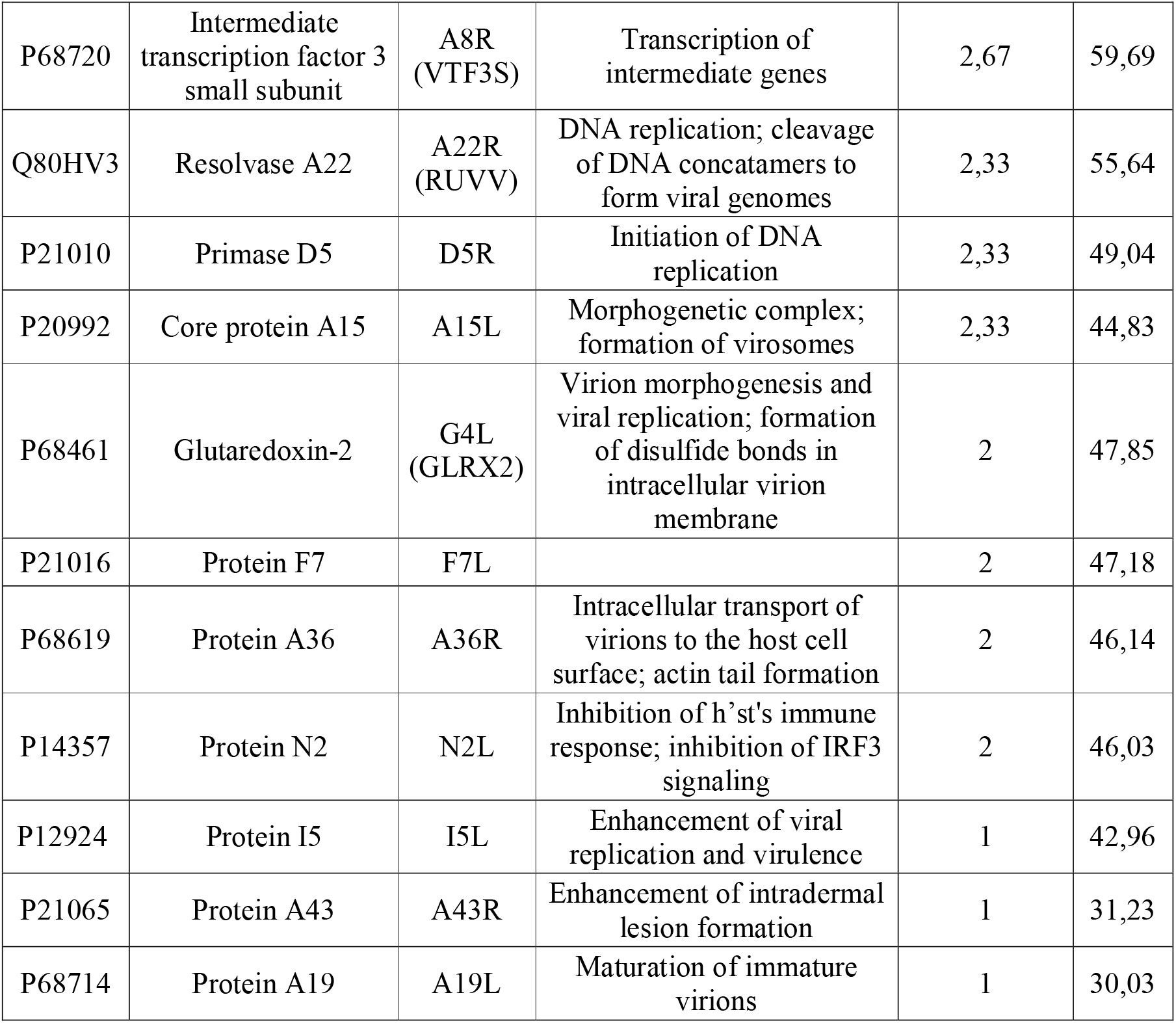
Immunoprecipitated viral proteins from purified IHD-J-ISG15 virions. Extracts from IHD-J-V5-ISG15 were immunoprecipitated using an V5 specific antibody and analyzed by LC-ESI-MS/MS. Table 3 lists: Protein ID (column A); protein name (column B); gene name (column C); protein function (column D); abundance, expressed as average peptides detected for each protein in the proteomic analysis of 3 biological replicates per condition (column E); statistical significance (-log P-value) (column F). Proteins are ordered by abundance. Protein extracts of purified IHD-J virions were immunoprecipitated as negative control.

**Figure 5.**
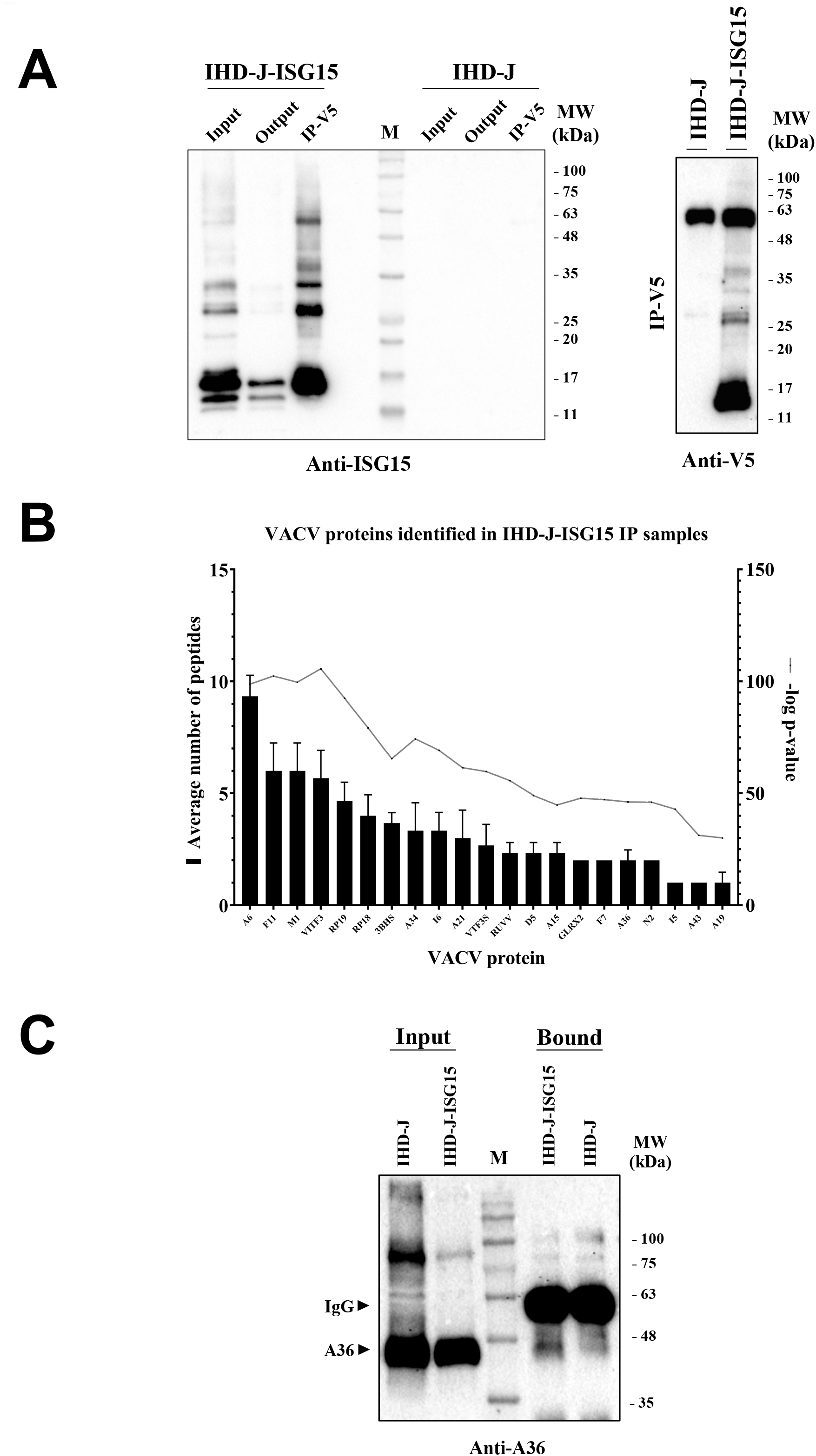
Several VACV proteins interact with ISG15. **(A) Immunoprecipitation of ISG15 and ISGylated proteins from IHD-J-ISG15 virions**. Sucrose gradient-purified IHD-J-ISG15 and IHD-J virions were processed, and V5-ISG15-conjugated viral proteins were subjected to immunoprecipitation using a V5 antibody. Left panel: The total protein extract before (Input) and after (Output) the immunoprecipitation assay, and the immunoprecipitated proteins (IP-V5) were fractionated by SDS-PAGE, and ISG15 and ISGylated proteins were analyzed by Western Blot using an ISG15-specific antibody. Right panel: The immunoprecipitated proteins were analyzed by Western Blot, using an anti-V5 specific antibody. Purified IHD-J virions were used as control. Molecular weights (MW) in kDa are indicated, based on protein standards (M). **(B) Identification of VACV proteins as potential targets of ISGylation**. Immunoprecipitated protein extracts from sucrose gradient-purified IHD-J-ISG15 and IHD-J virions were analyzed by LC-ESI-MS/MS to identify VACV proteins that interact with ISG15. The graph collects 21 proteins that were found exclusively in IHD-J-ISG15 IP samples, identified as potential ISG15 interactors. The average number of peptides and -log p-value from 3 biological replicates are represented for each protein. **(C) Validation of the VACV A36-ISG15 interaction by immunoprecipitation**. Purified IHD-J-ISG15 virions were processed, and V5-ISG15-conjugated viral proteins were subjected to immunoprecipitation using a V5 antibody. Thfe total protein extract (Input) and the immunoprecipitated protein extract (Bound) were fractionated by SDS-PAGE, and the interaction between VACV A36 and ISG15 was analyzed by Western Blot using an A36-specific antibody. Purified IHD-J virions were used as control. Molecular weights (MW) in kDa are indicated, based on protein standards (M).

Altogether, these results indicate that ISG15 modulates the proteome of VACV, what is expected to impact on VACV infection at different levels.

## Discussion

Poxviruses are the only known viruses that produce more than one infectious form during their infectious cycle. The distinct infective particles differ not only antigenically and structurally, but also in the way that they exit the cell (17, 19, 40). While MVs are released from the infected cell after cell lysis, the EVs bud directly from the plasma membrane without necessarily encouraging cellular death. Although *Variola virus* (VARV; the causal agent of smallpox) has been eradicated, poxviruses are currently back in the news after the recent outbreak of monkeypox (MPXV), a zoonotic *Orthopoxvirus* that also causes disease in humans, although it results in notably lower mortality rates compared with VARV (41). The possibility that, in the future, MPXV conquers the ecological niche once occupied by VARV exists; therefore, it is important to understand the host restriction factors that control the different *Poxvirus* dissemination mechanisms.

IFN is an essential factor for viral restriction, and although poxviruses encode multiple proteins that prevent its action, occasionally they are not able to counteract its effect completely (42). For example, during infection of bone marrow-derived macrophages (BMDM), VACV is susceptible to IFN treatment (29). ISG15 is one of the most highly expressed genes in response to IFN, and its direct and indirect antiviral action is well established (9). Recently, we have shown that ISGylation inhibits the production of exosomes (31), vesicles secreted to the extracellular environment, very similar to EVs. Considering that it has been discussed that EVs employ a mechanism similar to exosindicateomes (25), here, we hypothesized that, as is the case with exosomes, ISG15 may act as a sensor for viral dissemination determining which of the infectious forms of VACV should be generated. Based on this hypothesis, in this work we set out to study whether the ISG15/ISGylation system could regulate VACV cycle and the production of MVs and EVs which may have important implication on viral spread. We used the WR and IHD-J strains of VACV, and although we demonstrated that ISG15 is essential for the correct dissemination in both strains, the higher production of EVs by IHD-J (43, 44) made us focus our work on this strain. Our results clearly evidence that the absence of ISG15 is associated with a reduction in EV production, in line with a severe impairment in the production of comet-shaped plaques. However, although there is a clear defect in EV release in *Isg15-/-* cells, the production of viral proteins did not differ between *Isg15+/+* and *Isg15-/-* cells, as observed with A27, the classical marker for MV/EV, and F13, the major envelope protein of EV, that associates with Golgi membranes and is essential for EV formation (17). These results suggest that the absence of ISG15 does not affect the correct kinetics of viral gene expression (34). The interaction of F13 with the MV surface is a requirement for the association between MVs and wrapping membranes (45) and its absence leads to a dramatic reduction of viral spread and plaque size (46). As mentioned above, we did not observe any differences in the levels of F13 between genotypes, although we cannot rule out that the absence of ISG15 affects the proper function of F13 in this process. IHD-J is very efficient in TGN-mediated virion wrapping and the release of extracellular enveloped virus (44); however, as seen by TEM, intracellular virus particles were increased in IHD-J-infected *Isg15-/-* cells. Surprisingly, when we quantified the intracellular infectious virions by plaque assay, we did not observe any difference between genotypes, suggesting that perhaps many of the accumulated particles in *Isg15-/-* cells might be defective and, therefore, not infectious. In addition, as our data indicate a decrease of infectious EVs in infected *Isg15-/-* cells, we hypothesize that the ISG15/ISGylation system plays a regulatory role during virion egress.

This irregular viral spread observed in *Isg15-/-* MEF also associated with a quantitative difference in the protein signature of IHD-J virions (Table 1-2). The quantitative proteomic analysis of gradient-purified intracellular virus showed a general increase of cellular and viral proteins in virions grown in *Isg15-/-* MEF compared to those grown in *Isg15+/+* MEF (Table 1). The fact that many mitochondrial proteins were present in purified virions might indicate that these proteins were co-purified with the virions, mainly involved in oxidative phosphorylation, those levels were upregulated in purified from *Isg15-/-* MEF (Table 2). We have previously shown that ISG15 is necessary for proper oxidative phosphorylation, and that the absence of ISG15 results in the accumulation of mitochondrial proteins (29). However, given the large size of VACV and its complexity, it is possible that many of these proteins were engulfed within the virions during the morphogenetic process. Interestingly, in our preliminary proteomic study of virion-enriched cell extracts (not shown), we found the Ring Finger Protein 213 (RNF213) among the less abundant cellular proteins in *Isg15-/-* samples. This protein was described as a sensor of ISGylated proteins, which showed antimicrobial activity against bacteria and viruses, *in vitro* and *in vivo* (47). This observation suggests that the interaction between ISG15 and RNF213 might have a relevant role in the antiviral response against VACV.

Focusing on viral proteins present in both types of virions, we identified 58 proteins significantly upregulated, and only 5 significantly reduced (N1, A34, F11, D13 and IL8B), in virions purified from *Isg15-/-* MEF comparing with those from *Isg15+/+* MEF (Table 1). Our results regarding the proteins identified in virions are in line with previous proteomic analyses of purified virions (48-50). Interestingly, we identified several proteins involved in the formation of EVs and actin tails, such as B5, F13, A34 and F11. The protein B5 was the most upregulated protein in virions purified from *Isg15-/-* cells. B5 has been shown to be involved in IEV formation and actin tail polymerization (51). A33, A34, and B5 are critical for the efficient production of infectious EV and interactions among these proteins are important for their localization and incorporation into the outer extracellular virion membrane (52). Our proteomic analysis reported a reduction in the levels of A34 in virions purified from *Isg15-/-* MEF. The absence of A34 was linked to a strong decrease in the amounts of B5 present in EV (52); however, little is known about how a dysregulation in A34 levels, other than its complete absence, affects B5. Furthermore, the presence of B5 at the surface of the EV is required for actin tail formation after contact with the membrane of cells expressing A33 and A36. Our results indicate that, in the absence of ISG15, the release of EV and the formation of actin tails are impaired. Moreover, in our study, we detected increased levels of B5 in virions purified from *Isg15-/-*MEF. We hypothesize that the absence of ISG15 causes alterations in virion egress, what leads to an accumulation of B5-containing viral progeny. This would explain the higher levels of B5 in purified virions from *Isg15-/-* MEF, and a reduction in actin tail formation, since these B5-containing virions would not be released and, therefore, could not induce actin polymerization at the cell surface.

The protein F11, involved in EV formation and virus spread (37), was identified as the most downregulated protein in virions from *Isg15-/-* MEF. F11 modulates the cortical actin cytoskeleton and enhances the release of virions (36); therefore, a reduction in the levels of F11 is consistent with our hypothesis of impaired virion egress and our observations of decreased EV release. Interestingly, we identified F11 and A34 as proteins that interact with ISG15 (Table 3). Considering that ISG15 regulates host and pathogen protein dynamics, it is possible that its absence alters the localization and interaction of viral proteins involved in virion maturation, actin tail formation and EV formation (10, 33).

A26 was one of the viral proteins increased in virions grown in the absence of ISG15, as reported by the proteomic analysis of purified virions and a previous proteomic analysis with virion-enriched cell extracts. Moreover, an increase in A26 levels was detected by Western Blot analysis of infected *Isg15-/-* MEF (not shown). The A26 protein is present in MVs but absent in wrapped virions (53); therefore, the accumulation of intracellular viral particles reported in *Isg15-/-* MEF could explain these observations. Furthermore, such accumulation could also explain the increased levels of A26 in purified virions from *Isg15-/-* MEF, as the content of MV is expected to be higher than in virions purified from *Isg15+/+* MEF. In some *Orthopoxvirus* species, A26 participates in the attachment of MVs to the cell surface and the incorporation of MVs into A-type inclusions (ATI) (54). Those structures provide stability and protection to MVs ensuring a proper transmission between surrounding cells, and the presence of the entire AT1 protein (A25) is required for its formation (54). In VACV, the AT1 protein is truncated and does not aggregate into ATI; however, it still associates with and retains A26 (55). Several studies suggest that A26 mediates in the decision between virion wrapping and the formation of A-type inclusions, suggesting that A26 acts as a switch to enhance the production of MVs at the expense of EVs (53). A26 also enhances retrograde transport of MVs (55) and participates in cell attachment by interacting with laminin (56). Recently, Holley et al., in line with our studies, demonstrated that A26 expression downregulates EV production, pointing out that MV maturation is controlled by the abundance of A26 (57).

Our work shed light on a novel mechanism in which ISG15 is a host factor for the control of EV production, in line with an accumulation of A26, and higher levels of relevant proteins for VACV, such as B5, and F13, in virions purified from *Isg15-/-* MEF. Therefore, we wondered whether ISG15 regulates these proteins through ISGylation. It has been shown that, during viral infections, the adequate regulation of post-translational modifications (PTMs) on the same or nearby residues of the same protein, is important to perpetuate the infection (12). Thus, it is possible that the ISGylation of VACV proteins alters other PTMs required for their proper function. Our previous efforts to immunoprecipitate ISG15 and ISGylated proteins from purified virions were unsuccessful, so we decided to generate a recombinant virus expressing V5-tagged ISG15. ISG15 is conjugated to *de novo* synthesized proteins in a co-translational manner, what facilitates ISGylation of viral proteins during infection (58). However, VACV downregulates ISG15 expression and shuts off translation of host proteins (59) and considering that ISGylation occurs to de novo synthesized proteins, the detection of ISGylated proteins after VACV infection is very difficult, especially at late stages of infection. To overcome this drawback, the expression of recombinant ISG15 was controlled by an early/late viral promoter that allows the expression of ISG15 through the viral cycle. Thus, ISG15 was synthesized at the same time as viral proteins, what increased the efficiency in ISGylation of viral proteins. Immunoprecipitation of V5-tagged ISG15 from IHD-J-ISG15 virion protein extracts and the subsequent proteomic analysis identified 21 VACV proteins capable of interacting with ISG15. Among them, we identified envelope proteins, such as A36, A34 and F11; core proteins, like A15, A19, A6 and A21; proteins with enzymatic activity, such as F7, GLRX2 and 3BHS; proteins involved in replication and transcription of the viral genome (e.g., VITF3, VTF3S, RP18, RP19 and RUVV), and proteins involved in the antiviral response, such as N2 (60).

The protein A36 is predominantly associated with the outer IEV envelope and, after virion release, it accumulates in plasma membrane beneath the CEV, where it is necessary for actin tail formation (61). It was demonstrated that multiple interactions between IEV membrane proteins exist, which are relevant for IEV assembly and actin tail formation (17, 62). An efficient virus release required a tight tethering to the host cell through interactions mediated by viral envelope proteins (63). The detection of an additional band of high molecular weight (~ 85 kDa) in our Western Blot analysis of A36 with virion protein extracts could be the result of one of these interactions (Fig. 5C). Moreover, this band is clearly reduced in IHD-J-ISG15 extracts, indicating that perhaps the presence of ISG15 and its interaction (covalent or not) with A36 alters its association with other proteins. As the proteins A34 and A36 participate in actin tail formation, the fact that they appear as potential ISG15 targets suggests that an impairment of the interplay between ISG15 and these proteins in *Isg15-/-* cells could also contribute to the reduction in actin tails. In future work we will try to elucidate the biological significance of this interaction (perhaps ISGylation) in the context of VACV infection. Unfortunately, the A26 protein was not present among our immunoprecipitated candidates, indicating that the interaction between ISG15 and A26 is unlikely to occur.

The major finding of the present manuscript is that ISG15 is required for the production of EVs; however, this should not be interpreted as a proviral effect of ISG15 during VACV infection, as it has been described for other viruses, such as HCV or HBV (Reviewed in (9)). We hypothesize that the fact that ISG15 favors the production of EVs might facilitate the early detection of these particles by the immune system and the rapid stimulation of immune responses, considering that EVs are susceptible to antibody-mediated neutralization (64-66). Although our hypothesis is entirely speculative, similar mechanisms have been observed during bacterial infections. It has been shown that the production of bacteria-derived extracellular vesicles are a vehicle for spreading of pathogen-associated molecular patterns (PAMPs), that stimulate the host’s innate immune response through the activation of pattern-recognition receptors (PRR), Stimulator of Interferon Genes (STING) and inflammasome signaling pathways (67).

Many questions remain open to the fact that poxviruses produce two different infectious forms with different characteristics. Is this advantageous? And, if so, why the percentage of EVs produced is so low compared to the MVs? Is it more advantageous to produce MVs in large quantities and wait for the cell to lyse or does it make sense to release virions into the extracellular medium as soon as possible? Regarding to the pathogenicity, it has been described that MVs promote direct cell-to-cell spread whereas EV budding enables dissemination from host-to-host and spread across long distances to infect distant tissues within the host (19). IHD-J-infected mice showed reduced mortality compared with WR-infected mice (43), suggesting that higher EV production and dissemination within the host does not significantly influence virulence; the WR strain, however, is highly pathogenic despite its low EV production [54]. Regarding VARV, in several strains the increased comet formation phenotypes and the subsequent EV production is associated with geographical and phylogenetic origin, but not with increased pathogenicity (68). With the current MPXV outbreak, the threat of its expansion and the probability of appearance of new mutants that could occupy the ecological niche left vacant by VARV, we consider that is timely and relevant to investigate the aspects of host–pathogen interactions that control *Poxvirus* transmission.

In summary, our results suggest that ISG15 is a novel host factor for the modulation of VACV egress and dissemination. In this work, we demonstrate that: i) ISG15 regulates EV production and the formation of comet-shaped plaques; ii) ISG15 regulates actin tail formation; iii) ISG15 has an impact on the viral proteome composition; iv) ISG15 interacts with several VACV proteins (e.g., A36), although the outcome of these interactions remains to be elucidated. Altogether, our work highlights an essential role for ISG15/ISGylation in viral infection and cell-to-cell spread and points out the relevance of the comprehension of ISG15-mediated antiviral responses, what might lead to the development of effective therapies against relevant human pathogens.

## MATERIALS AND METHODS

### Cells and viruses

Immortalized *Isg15+/+* and *Isg15-/-* MEF (kindly provided by K.P. Knobeloch) (69) and NIH/3T3 (ATCC^®^ CRL-1658^™^) cells were cultured in Dulbecco’s modified Eagle’s medium (DMEM) supplemented with 10% fetal calf serum (FCS). BSC40 (ATCC^®^ CRL-2761^™^) cells were cultured in DMEM supplemented with 5% FCS. FCS used in all experiments was heat inactivated (56°C, 30 min) prior to use. NIH/3T3 were transduced with lentiviral particles containing specific shRNAs for ISG15, obtaining 3T3-L1 and 3T3-L2, or a scramble shRNA, obtaining 3T3-SC. To generate viral stocks, the International Health Department-J (IHD-J) strain of vaccinia virus (VACV) (kindly provided by Dolores Rodriguez), the IHD-J-ISG15 recombinant virus, and the Western Reserve (WR) strain of VACV, were grown in BSC40 cells (0.01 PFU/cell; 48 h) and purified by centrifugation through a sucrose cushion followed by a sucrose gradient, as previously described (70) Virus aliquots were stored at -80ºC.

### Virus titration

Viral stocks as well as samples of intracellular or extracellular virus were titrated by plaque assay as previously described (71), with slight modifications: DMEM-2% FCS containing 1% low temperature melting agarose was used as overlay medium for the IHD-J strain. Intracellular virus samples were subjected to 3 cycles of freezing-thawing to allow for the cell lysis. Extracellular virus samples were titrated freshly after centrifugation during 2 min at 250 ^x^g.

### Comet-shaped plaque formation assays

Cell monolayers in 6 well plates were infected with approximately 0.0001 PFU/cell of IHD-J or WR per well in DMEM. After 1 hour of incubation at 37ºC and 5% CO2, cells were washed and incubated with DMEM supplemented with 2% FCS, for additional 48 (for IHD-J) or 72 h (for WR). Plaques were visualized after fixation and staining with 0.1% crystal violet in 10% formaldehyde in PBS.

### Immunofluorescence

Immortalized *Isg15+/+* or *Isg15-/-* MEF were grown on 12-mm-diameter glass coverslips in DMEM–10% FCS to a confluence of 40% to 50% and mock infected or infected at a multiplicity of infection (MOI) of 2 PFU/cell with IHD-J. At the indicated times post infection, cells were washed with phosphate-buffered saline (PBS), fixed with 4% paraformaldehyde, permeabilized with 0.25% Triton X-100 in PBS for 30 min, and blocked in PBS with 10% FCS for 45 min at room temperature. Specific monoclonal antibodies for the VACV A27 protein (1:1000; kindly provided by Mariano Esteban), F13 protein (1:500; kindly provided by Rafael Blasco) were used as primary antibodies. Alexa Fluor 488-conjugated mouse (for A27) or 594-conjugated rat (for F13) IgG antibodies (Invitrogen) were used as secondary antibodies. Cell nuclei were stained with 4′,6-diamidino-2-phenylindole (DAPI) (Sigma) (1:200). F-actin was stained with rhodamine-conjugated phalloidin (Molecular Probes). Confocal microscopy was performed using a Leica SP8 laser scanning microscope, and images were collected and processed with LAS X software (Leica, Wetzlar, Germany).

### Electron microscopy

Monolayers of *Isg15+/+* or *Isg15-/-* immortalized MEFs were infected at an MOI of 2 PFU/cell with the strain IHD-J of VACV. At 9 hpi, when the cytopathic effect was evident, the supernatant was removed and the cells were fixed with a solution of 2.5% glutaraldehyde containing 1% tannic acid, 0.4 M HEPES in PBS. After fixation, cells were carefully scraped, centrifuged to eliminate the fixative, and processed for embedding in the epoxy-resin EML-812 as previously described (72). Electron micrographs were taken using a transmission electron microscope (JEOL JEM-1011; Centro Nacional de Biotecnología, Spain) equipped with a ES1000W Erlangshen charge-coupled-device (CCD) camera (Gatan Inc.) at an acceleration voltage of 40 to 100[kV. Fifty low-magnification micrographs were analyzed per genotype. Cells which can be visualized in its integrity were considered for the analysis. Non-infected cells were excluded from the analysis. The number of intracellular viral particles per cell was quantified, with more than 110 cells analyzed per genotype.

### Protein analysis by Western Blotting

Immortalized *Isg15+/+* or *Isg15-/-* MEF, NIH-3T3 cells and lentivirus-transduced NIH-3T3 cells, or purified virions were collected, and lysates were obtained by solubilizing cells in SDS-Laemmli sample buffer supplemented with 100 mM DTT. Cell lysates were boiled for 5 min, resolved by sodium dodecyl sulfate-polyacrylamide gel electrophoresis (SDS-PAGE) with Laemmli running buffer and transferred to PVDF membranes (Merck-Millipore) in a Trans-Blot^®^ SD Semi-Dry Transfer Cell (Bio-Rad), following the manufacturer’s recommendations. Membranes were blocked with 5% skim milk in PBS containing 0.1% Tween-20 (PBS-T) and incubated with the corresponding primary antibodies as indicated in the figure legends (A27 and A4 kindly provided by Mariano Esteban; F13 and A36 kindly provided by Rafael Blasco and A26 kindly provided by Wen Chang) in 0.5% skim milk in PBS-T. Membranes were then washed with PBS-T and incubated with anti-rabbit or anti-mouse peroxidase-labeled antibodies (1:10000, Sigma-Aldrich). After extensive washing with PBS-T, the immune complexes were detected using Clarity Western ECL blotting substrate (Bio-Rad) and a ChemiDoc XRS^+^ System (Bio-Rad), according to the manufacturer’s instructions. Protein quantity was analyzed using Fiji software (73).

### Generation of recombinant IHD-J-ISG15

Monolayers of BSC-40 cells were infected with parental IHD-J at an MOI of 0.01 PFU/cell and transfected 1 hour later with 10 μg of DNA of plasmid pCyA-ISG15GG (pCyA was kindly provided by Mariano Esteban) using LT-1 (Mirus Bio) according to manufacturer’s recommendations. The gene ISG15GG was fused to a V5 tag sequence at its N-terminal domain to facilitate its detection. At 48 hpi, the cells were harvested, lysed by freeze-thaw cycling, sonicated, and used for recombinant-virus screening. Recombinant IHD-J containing the 513-bp DNA fragment encoding the ISG15-V5 fusion protein and transiently co-expressing the β-Gal reporter gene (IHD-J-ISG15, X-Gal^+^) were selected by three consecutive rounds of plaque purification in Neutral Red (Sigma, Cat. No. N2889) and X-Gal (5-bromo-4-chloro-3-indolyl-β-d-galactopyranoside, 40 mg/mL)-stained BSC-40 cells. In the following purification steps, recombinant IHD-J containing the ISG15 gene and lacking the β-Gal gene (deleted by homologous recombination between the TK left arm and the short TK left arm repeat flanking the reporter gene) (IHD-J-ISG15, X-Gal^−^), were isolated by three additional consecutive rounds of plaque purification in X-Gal-stained (40 mg/mL) BSC40 cells, screening for non-stained viral plaques. In each round of purification, the isolated plaques were expanded in BSC-40 cells for 24 hours, and the viruses obtained were used for the next purification round. To test the identity and purity of the recombinant virus IHD-J-ISG15, DNA was extracted from infected cells and subjected to PCR amplification using the oligonucleotides TK-L (5′-TGATTAGTTTGATGCGATTC-3′) and TK-R (5′-TGTCCTTGATACGGCAG-3′), annealing in the TK flanking sequences. The resulting PCR product was analyzed by sequencing. ISG15 expression was analyzed by Western Blot using anti-V5 antibody (Invitrogen, Cat. No. R960-25).

### Statistical analysis

One-tailed or two-tailed unpaired Student’s *t* tests were used to analyze the differences in mean values between groups. All results were expressed as means ± standard deviation; *P* values of <0.05 were considered significant.

### LC-MS/MS of purified virions

*Isg15+/+* or *Isg15-/-* MEF were infected with IHD-J (MOI 0.01, 48 h), scraped in culture medium, pelleted and resuspended in 10□mM Tris pH 9.0. Cells were disrupted and vigorous vortexing for 90□s. After sonication for 3□min, cell debris were pelleted at 1,000□×□*g* and 4□°C for 5□min. Purification of mature virions was done by rate-zonal sucrose gradient centrifugation in sterile SW 28 centrifuge tubes (PMID: 18265124). For this purpose, the supernatant was centrifuged through a 20% sucrose cushion at 32,900□×□*g* and 4□°C for 80□min and the virus pellet was resuspended in 1□ml of 10□mM Tris pH 9.0. The virus suspension was sonicated for 1□min, layered on a 20% to 45% continuous sucrose gradient and centrifuged at 26,000□×□*g* and 4□°C for 50□min. The virus bands were pooled and concentrated at 32,900□×□*g* and 4□°C for 60□min. The pellet was resuspended in 1□ml of 10□mM Tris pH 9.0 and stored in aliquots at −80□°C. Purified virions were processed for LC-MS/MS analysis. ***In-solution Digestion***. Protein samples were individually digested with trypsin using a standard protocol. Briefly, 20 µg of protein of each sample were resuspended and denatured in 20 µL of 7 M urea, 2 M thiourea, 100 mM TEAB (triethylammonium bicarbonate), reduced with 2 µL of 50 mM Tris 2-carboxyethyl phosphine (TCEP) (AB SCIEX, Foster City, CA, USA), pH 8.0, at 37 °C for 60 min and followed by cysteine-blocking reagent chloroacetamide (CAA). Samples were diluted up to 60 µL with 50 mM TEAB to reduce the concentration of urea. One µg of sequence grade-modified trypsin (Pierce) was added to each sample (ratio 1:20 enzyme:sample), which were then incubated at 37 °C overnight on a shaker. After digestion, samples were dried in a SpeedVac (Thermo Scientific, Waltham, MA, USA). ***Tagging with TMT 6plex***^***TM***^ ***reagent***. The resulting tryptic peptides were subsequently labelled using TMT-6plex Isobaric Mass Tagging Kit (Thermo Scientific, Rockford, IL, USA) according to the manufacturer’s instructions as follows: 126: WT-R1; 127: WT-R2; 128: WT-R3; 129: KO-R1; 130: KO-R2; 131: KO-R3). After labelling, the samples were pooled, evaporated to dryness, and stored at -20°C until the LC−MS analysis. Three biological replicates of each condition were analyzed. ***Liquid chromatography and mass spectrometry analysis (LC-ESI-MS/MS)***. Before MS analysis, we determined the amount of peptide in the combined sample by Qubit™ Fluorometric Quantitation (Thermo Fisher Scientific). A 1 µg aliquot of each fraction was subjected to 1D-nano LC ESI-MS/MS (Liquid Chromatography Electrospray Ionization Tandem Mass Spectrometric) analysis using an Ultimate 3000 nano HPLC system (Thermo Fisher Scientific) coupled online to an Orbitrap Exploris 240 mass spectrometer (Thermo Fisher Scientific). Peptides were eluted onto a 50□cm□×□75 μm Easy□spray PepMap C18 analytical column at 45°C and were separated at a flow rate of 300□nL/min using a 120 min gradient ranging from 2 % to 35 % mobile phase B (mobile phase A: 0.1% formic acid (FA); mobile phase B: 80 % acetonitrile (ACN) in 0.1% FA). The loading solvent was 2 % ACN) in 0.1 % FA and injection volume was 5 µL. Data acquisition was performed using a data-dependent top-20 method, in full scan positive mode, scanning 375 to 1200 m/z. Survey scans were acquired at a resolution of 60,000 at m/z 200, with Normalized Automatic Gain Control (AGC) target (%) of 300 and a maximum injection time (IT) in AUTO. The top 20 most intense ions from each MS1 scan were selected and fragmented via Higher-energy collisional dissociation (HCD). Resolution for HCD spectra was set to 45,000 at m/z 200, with AGC target of 100 and a maximum ion injection time in AUTO. Isolation of precursors was performed with a window of 0.7 m/z, exclusion duration (s) of 45 and the HCD collision energy was 30. Precursor ions with single, unassigned, or six and higher charge states from fragmentation selection were excluded. Raw instrument files were converted to MGF files and MS/MS spectra searched using OMSSA 2.1.9, X! TANDEM 2013.02.01.1, Myrimatch 2.2.140 and MS-GF+ (Beta v10072) against a composite target/decoy database built from the *Mus musculus* reference proteome sequences and the Vaccinia virus (strain Western Reserve) proteome downloaded from UniprotKB. Search engines were configured to match potential peptide candidates with mass error tolerance of 25 ppm and fragment ion tolerance of 0.02 Da, allowing for up to two missed tryptic cleavage sites and a maximum isotope error (13C) of 1, considering fixed carbamidomethyl modification of cysteine and variable oxidation of methionine, pyroglutamic acid from glutamine or glutamic acid at the peptide N-terminus, and modification of lysine and peptide N-terminus with TMT 6-plex reagents. Score distribution models were used to compute peptide-spectrum match p-values, and spectra recovered by a false discovery rate (FDR) _ 0.01 (peptide-level) filter were selected for quantitative analysis. Approximately 5% of the signals with the lowest quality were removed prior to further analysis. Differential regulation was measured using linear models, and statistical significance was measured using q-values (FDR). All analyses were conducted using software from Proteobotics (Madrid, Spain).

### Immunoprecipitation assay

Sucrose gradient-purified intracellular virions of IHD-J and IHD-J-ISG15 strains, previously sonicated, were resuspended in a 50 mM Tris-HCl (pH 7.5) buffer, containing 150 mM NaCl, 1% NP40 detergent and supplemented with a protease inhibitor cocktail mini tablet (complete from Roche) and a phosphatase inhibitor mini tablet (Pierce from Thermo Scientific) according to the manufacturer’s instructions. The virus extract was vortexed and incubated on ice for 15 minutes. The amount of protein in the samples was determined by the Bradford protein method. For immunoprecipitation: 70µg of protein extract were incubated with the anti-V5 antibody (diluted 1:50) overnight at 4 °C, then protein G-Sepharose beads were added to the mixture and the sample was further incubated for 2 hours at 4 °C. The beads were washed four times with a 50 mM Tris-HCl (pH 7.5) buffer containing 150 mM NaCl and 0.1% NP40 detergent, and finally washed once with PBS. The immunoprecipitated proteins were eluted with SDS sample buffer and analyzed by immunoblot.

### Protein identification of immunoprecipitated extract

Protein eluted from the immunoprecipitation assay (three biological replicates per condition) were individually loaded and concentrated in a 12% SDS-PAGE gel. Each sample was divided in 4-5 bands, and these were digested with trypsin, using an automatic robot Opentrons (www.opentrons.com/contact). In all cases, digestion was performed according to a protocol described by Shevchenko et al **(74)**. In summary, gel plugs were washed with 50 mM ammonium bicarbonate and samples reduced with 10 mM DTT. Alkylation was carried out with 10 mM chloroacetamide at room temperature before adding recombinant sequencing-grade trypsin (0.1 μg, Promega). Digestion took place at 37º C for 18 h. Following digestion, peptides were extracted, pooled, dried by speed-vac centrifugation, and stored at -20ºC until needed.

### LC-ESI-MS/MS analysis and database searching

Nano LC ESI-MS/MS analysis was performed using a Thermo Ultimate 3000 nanoHPLC coupled to a Thermo Orbitrap Exploris 240 mass spectrometer (ThermoFisher Scientific, San José, California, USA). The analytical column used was an Easy-spray PepMap RSLC C18 reversed phase column 75 µm × 50 cm, 2 µm particle size. The trap column was an Acclaim PepMap 100, 5 µm particle diameter, 100 Å pore size, switched on-line with the analytical column. The loading pump delivered a solution of 0.1% formic acid in 98% water / 2% acetonitrile (Merck, Germany) at 10 µL/min. The nanopump provided a flow-rate of 250 nL/min and was operated under gradient elution conditions, using 0.1% formic acid (Fluka, Buchs, Switzerland) in water as mobile phase A, and 0.1% formic acid in 80% acetonitrile as mobile phase B. Gradient elution was performed according the following scheme: isocratic conditions of 94% A: 6% B for two minutes, a linear increase to 35% B in 40 min, a linear increase to 95% B in one minute, isocratic conditions of 95% B for four minutes and return to initial conditions in 1 min. Injection volume was 5 µL. The LC system was coupled via an Easy-sprayTM nanospray source to the mass spectrometer. Automatic data-dependent acquisition using dynamic exclusion allowed obtaining both full scan (m/z 350-1250) MS spectra followed by tandem MS HCD spectra of the 20 most abundant ions. MS and MS/MS data were used to search against a composed database containing *Chlorocebus sabaeus* and *Vaccinia virus* protein sequences (downloaded from UniProtKB, june 2021). The database also contained a short list of common laboratory contaminants downloaded from CRAP Database (www.crapome.org). Searches were done using a licensed version of Peaks v.7.5 and search parameters were set as follows: carbamidomethyl cysteine as fixed modification and acetyl (protein N-term), NQ deamidation, -GG at Lys residues, Glu to pyro-glutamic and oxidized methionine as variable ones. Peptide mass tolerance was set at 10 ppm and 0.02 Da for MS and MS/MS spectra, respectively, and 2 missed cleavages were allowed. False Discovery Rate was set at 1% at peptide level. Only those proteins with at least one unique peptide were considered.

## Supporting information

Supplementary Figure 1. Lentivirus-mediated knockdown of ISG15 in NIH-3T3 cells blocks the formation of comet-like plaques during IHD-J infection. To

