## Supplementary figures and images for "ISG15 is required for the dissemination of *Vaccinia virus* extracellular virions"

### Supplementary Figure 1. Lentivirus-mediated knockdown of ISG15 in NIH-3T3 cells blocks the formation of comet-like plaques during IHD-J infection. To

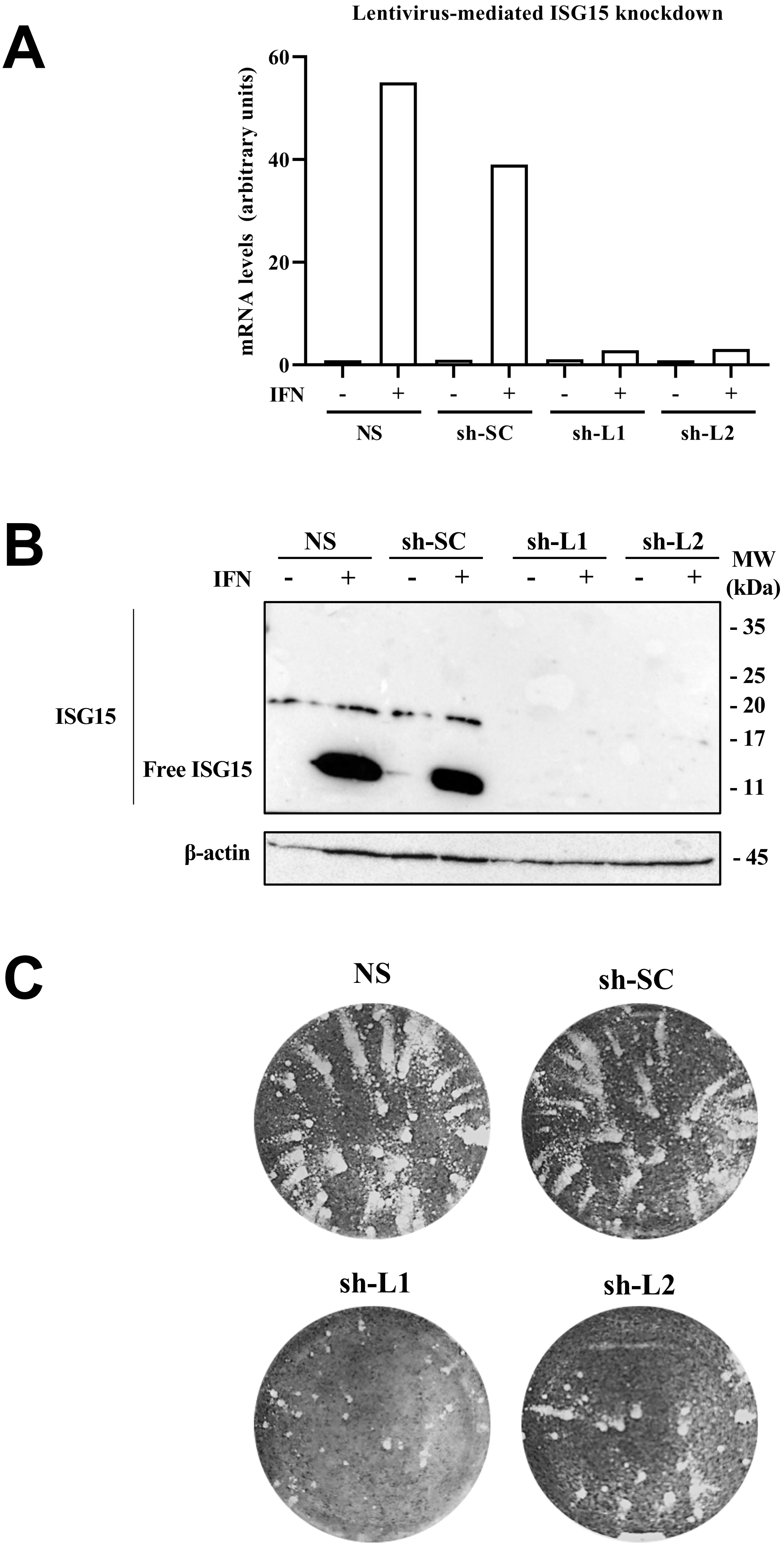

#### Figure S2

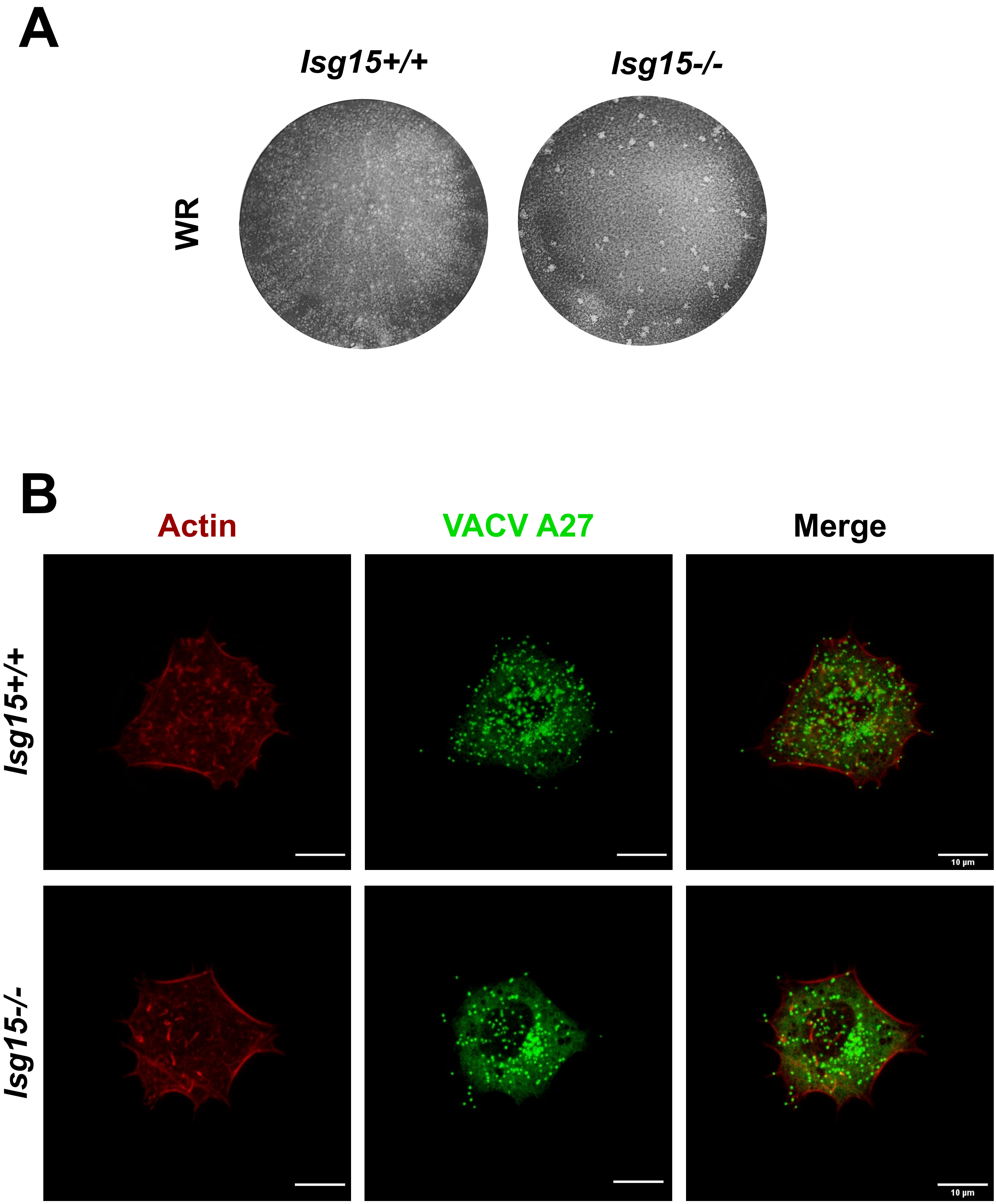

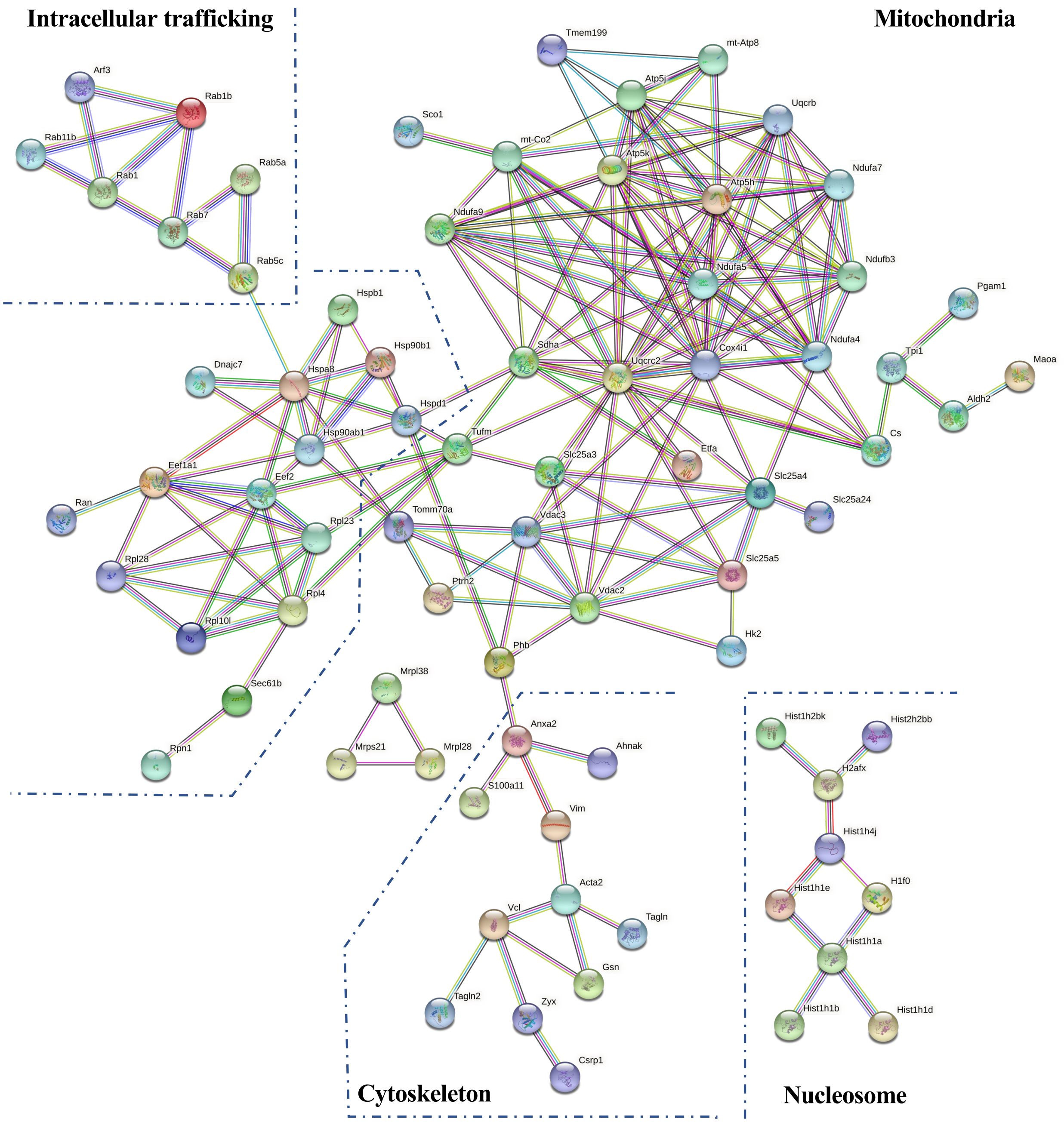
